# On why and how to encode probability distributions on graph representations of omics data: enhancing predictive tasks and knowledge discovery

**DOI:** 10.64898/2026.02.19.706756

**Authors:** Daniel M. Gonçalves, André Patrício, Rafael S. Costa, Rui Henriques

## Abstract

The growing availability and complexity of omics data have driven the development of specialized algorithms for modeling molecular systems. Although graph-based learning methods effectively represent biological interactions, they often neglect the statistical information embedded in node and edge annotations. To address this limitation, we propose a novel graph-based framework that integrates structured statistical distributions into nodes and edges, capturing probabilistic characteristics of molecular relationships. We evaluate the proposed approach on omics datasets from five cancer types across multiple clinical outcomes, including survivability and primary tumor site. Results demonstrate predictive performance comparable to established machine learning baselines. Beyond prediction, the statistically enriched graph representations enable the identification and characterization of regulatory modules associated with clinical outcomes, enhancing biological interpretability. These findings suggest that incorporating structured statistical information into graph representations provides a competitive and interpretable framework for predictive modeling and knowledge discovery in complex diseases.

## 1 Introduction

The rise in omics data sources has accelerated research efforts to model the regulatory mechanisms underlying complex diseases such as cancer [1, 2]. The high volume of profiled molecular features and their intricate interdependencies result in high levels of complexity that pose challenges for both descriptive and predictive modeling tasks. To overcome these challenges, specialized algorithms are necessary to effectively represent and extract information from these data sources [3, 4].

Conventional analysis methods are unable to model the rich nature of the interdependencies among biological entities, such as RNA molecules or proteins, often disregarding holistic perspectives on collective functionality [5, 6]. For this reason, the focus has been directed to the analysis of biological entity groups and their relations, rather than weighted cumulative views from isolated entities [7, 8], summarizing statistics or correlation analysis [9, 10].

Graphs are a well-known option for representing interactions between entities, offering the ability to represent complex data structures using node, layer, and edge constructs. Network-based approaches to omics data modeling offer the expressivity needed to represent complex biological systems, typically employing simple graphs, hypergraphs, or networks with unary, weighted, or multivariate annotations, which can include embeddings of arbitrarily high complexity [11]. As a result, graphs are frequently used in omics research, particularly gene expression data analysis [12, 13, 14, 15, 16]. Extensions have been proposed to represent multiple omic layers to capture the complex molecular interplay implicit in the central dogma [17, 18, 19, 20]. Classic network-based methods often assume that all direct neighbors of a known gene are phenotype-associated or directly co-regulated, overlooking complex long-chain interactions [21]. While broader perspectives have been explored, the high interconnectivity of biological networks poses challenges [22, 23].

More advanced methods aimed at addressing these challenges have been proposed. One approach, PARADIGM, models gene interactions as factor graphs to infer pathway alterations, improving robustness in breast cancer and glioblastoma, but remains limited to individual pathways [24]. SNF (Similarity Network Fusion) constructs patient similarity networks for each omic platform and fuses them iteratively, aiding in cancer subtyping and survival prediction [25, 26]. Other graph-based approaches use semi-supervised learning on multi-omics data to classify clinical outcomes by integrating multiple graphs and curated genomic knowledge through similarity matrices, capturing gene interactions from regulatory networks [18].

This work proposes a novel class of graph-based representations for the probabilistic modeling and subsequent predictive analysis of multi-omics data through an enhanced contrast analysis. We hypothesize that graph-based representations sensitive to the statistical distribution of relevant features of complex biological systems, as well as their corresponding impact on a given phenotype, are critical for enhanced prediction and knowledge discovery. As an alternative to simple, summarized views, the proposed methodology leverages graphs’ natural suitability as complex data structures capable of representing biological entities and their interactions through full probability mass and density distributions.

The major contributions of this work are:

1. A novel graph-based representation that encodes probability distributions on nodes and edges, enabling improved descriptive analysis of omics data;
2. New predictive models that effectively leverage the proposed graph representations, demonstrating robust learning capabilities even in datasets with limited samples and highly imbalanced target distributions;
3. Comprehensive empirical validation across multiple datasets from The Cancer Genome Atlas (TCGA), covering diverse cancer types, omic layers, and predictive tasks, showcasing the method’s practical applicability.

Section 2 describes the proposed methodology, while section 3 details the experimental setup. Section 4 presents the results from the experiments conducted on omics data obtained from TCGA platform, including 5 different cancer types [27]. Section 5 provides a more in-depth discussion of these results, including aspects of knowledge discovery and biological interpretability. Finally, conclusions are drawn in section 6.

## 2 Methods

The proposed methodology devises an effective graph representation of multi-omics data able to encode the rich interplay between molecular entities. In classical graph representations, *G* = (*V, E*), *nodes, V*, encode entities, such as genes, RNA molecules, proteins, or metabolites, and *edges, E*: *V* ×*V* → *𝒴*, encode relationships between the entities, where 𝒴 can be a unary association (i.e., {0, 1}), a quantity R, or a multivariate embedding. Although graphs offer suitable structures to capture interactions between biological entities, they generally disregard the inherent stochastic nature of these interdependencies. In contrast with classical representations, we explore the inherent expressivity of graphs to model the statistical distribution of molecular features and relationships through probability functions on nodes and edges. The proposed methodology is outlined in Figure 1.

**Figure 1:**
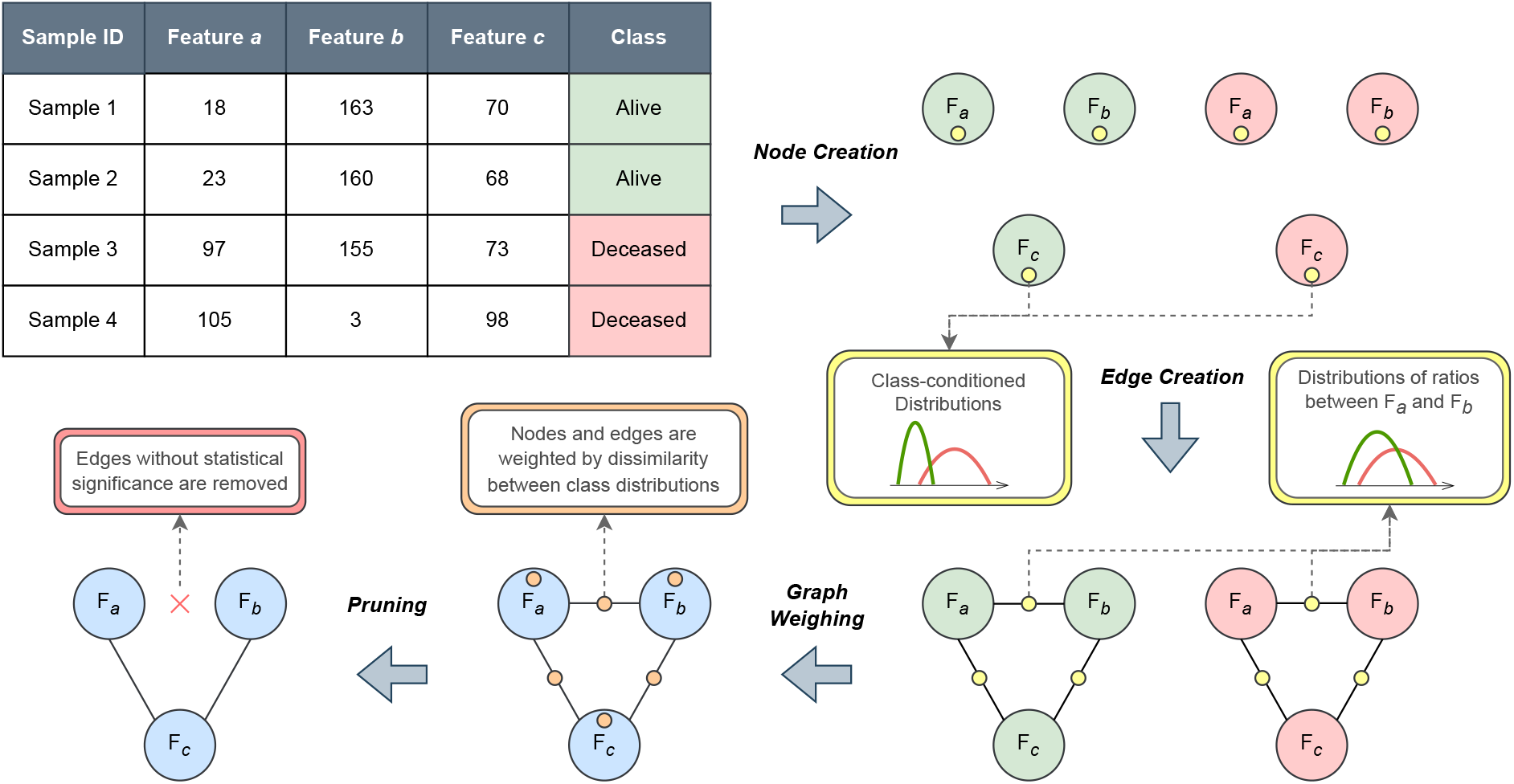
Overview of the graph construction methodology. During the *node and edge creation* steps, class-conditional probability functions (density or mass) are annotated on nodes and edges, represented by yellow dots. In the *graph weighting* step, weights are assigned based on the dissimilarity between class-conditional distributions at each node or edge, represented by orange dots. Finally, a *pruning step* removes edges that do not meet a statistical significance threshold.

In essence, each feature in a dataset, such as the expression of a given gene or concentration of a given metabolite, is mapped into a node in the graph. Each node stores the empirical distribution function of an associated feature, *V*: Ω → *f*_*X*_, which can be Probability Mass Functions (PMFs) for categorical features (e.g., genomic variation) or Probability Density Functions (PDFs) for numerical features (e.g., molecular concentration). In unsupervised settings, empirical distributions are inferred from all available measurements. In contrast, in supervised settings, class-conditional empirical distributions, *V*: Ω → {*f*_*X*|*c*_}_*c*∈𝒞_, one for each target *c* ∈ 𝒞, are approximated for each feature (node). The number of distribution functions per node essentially depends on the cardinality of discrete targets in classification tasks or numerical ranges in regression tasks. The graph creation process begins by pairing nodes to form potential edges. The edges also express (class-conditional) empirical probability functions, 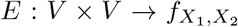. However, they reflect the contrast between the distributions of the associated nodes.

Each structure, node or edge, stores the class-conditional distributions and additional related statistics relevant for weighing and pruning purposes. These elements, which model the distribution of molecular features and their stochastic relationships, can be used directly for descriptive ends (knowledge acquisition) or, when conditioned to outcomes of interest, further guide predictive tasks.

### 2.1 Graph Generation

Let *X* be a matrix gathering data from an omics platform, where *i* ∈ 1, …, *n* indexes observations, *X*^*i*^, and *j* ∈ 1, …, *m* indexes features. Features generally correspond to measurements on biological entities (e.g., gene expression). Observations can be annotated with targets. Targets are denoted by **y**, where *y*_*i*_ corresponds to a target *c* ∈ 𝒞 (e.g., “alive” or “deceased”) for the *i*^*th*^ observation. Additionally, let *α* denote the reference significance-level threshold used in statistical tests.

#### Edges

To construct the target graph from the matrix *X*, edge formation is determined by pairwise comparisons between features. For instance, paired differences or ratios can be explored to produce new features, as well as the corresponding statistical distributions, able to encode the associative strength between features (nodes).

For two features *a* and *b* (*a* ≠ *b*), let their log-ratio transform be the pairing criterion,

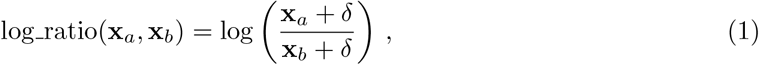

where *δ* is a (small) positive constant. The resulting values are stored in an *n*-dimensional vector, **v**, providing a unified representation of the relationship between the two features.

In unsupervised contexts, an empirical probability function is approximated from **v**, which is used as an annotation in the edge connecting nodes *a* and *b*.

In supervised contexts, a splitting step of measurements **v** by class is further performed for approximating |𝒞| class-conditional empirical probability functions to annotate the same edge.

For each pair of targets (*y*_*a*_, *y*_*b*_), where *a* ≠ *b*, the empirical distributions of **v** for samples in *y*_*a*_ and *y*_*b*_, denoted as 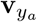 and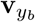, are compared using a statistical test, such as the Kolmogorov-Smirnov (KS) test [28],

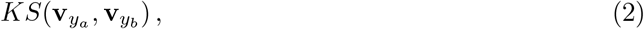

for assessing the equality of the empirical probability functions 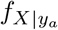 and 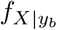 inferred from 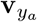 and 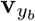. The test returns the corresponding *p*-value and score (e.g., KS test statistic).

In multiclass settings, |𝒞 | *>* 2, multiple *p*-values and scores are acquired,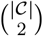. In this context, specific estimators of interest, including the minimum *p*-value or the maximum KS statistic, can be applied for retrieving a single summary statistic of the discriminative power of the precomputed ratios.

Complementary to probability functions, the introduced statistics of contrast can be used to further produce weight annotations on edges, *V* × *V* → ℝ. The computed *p*-values can be compared against a predefined statistical significance threshold, *α*, for filtering edges, ensuring that only those interactions that capture meaningful differences between targets are retained. This pruning step helps preserve the edges that are more informative for predictive tasks, further aiding the graph exploration for knowledge discovery ends.

#### Nodes

The node creation process follows a similar but simpler process. Given an individual feature **x**_*j*_, the available measurements are used to assign empirical probability functions and weights to the corresponding node. In supervised contexts, measurements are divided according to targets to fit class-conditional empirical probability distributions, 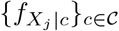. In addition, the predictive power of a node can be retrieved through scores such as the Kolmogorov-Smirnov (KS) statistic. For a given pair of classes, the similarity of the corresponding empirical distribution functions can be tested to produce a KS statistic. When |𝒞| *>* 2, an estimator, such as the maximum KS statistic from all pairs of classes, can be used to further annotate nodes with quantities.

### 2.2 Prediction

In the context of predictive tasks, the likelihood of each class is estimated for each testing instance by constructing instance-specific graphs. Given a testing instance, a new dedicated graph is created by testing the empirical probability functions stored on the nodes and/or edges of the original graph (computed during training time) at specific testing values. In other words, the predictor first computes feature likelihoods by assessing the class-conditional probability functions on all nodes at the testing features,

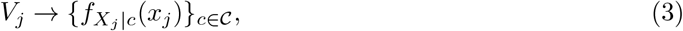

as well as ratio likelihoods by precomputing the log ratios between the features of the testing instance and assessing the class-conditional probability functions on the edges at these values,

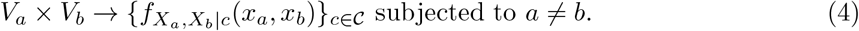

These estimates are used to create an instance-specific graph, *G* = (*V, E*), with multivariate annotations, ℝ^|𝒞|^, on nodes in *V* and/or edges in *E*.

To avoid overfitting risks posed by the precomputed empirical distributions, Kernel Density Estimation (KDE) is applied to this end [29, 30]. As default, KDE is parameterized with a Gaussian kernel with Scott’s rule as the thumb rule for bandwidth selection [31].

Given a specific instance *i*, and its corresponding testing graph, *G*_*i*_, the goal is to predict the class label ŷ_*i*_. The predictive process involves the following steps.

First, for each target *c* ∈ 𝒞, a likelihood score *ℓ*_*c*_ is computed based on the aggregation of node and edge weights in *G*. Formally, the likelihood for target *c* is given by:

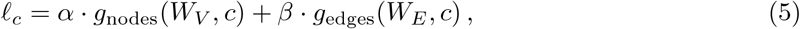

where *W*_*V*_ and *W*_*E*_ are the sets of node and edge weights, respectively; *g*_nodes_ and *g*_edges_ are aggregation functions that summarize node and edge contributions for target *c*. For simplicity, these functions are linear by default (i.e., weighted averages); *α* and *β* are hyperparameters controlling the relative contributions of node and edge information.

The acquired likelihood scores are transformed into probabilities using the softmax function,

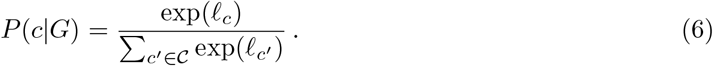

The predicted target ŷ is the one with the highest posterior probability,

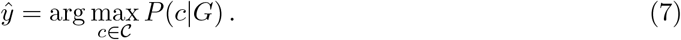

The optimization of the introduced graph-based predictors is guided by four major parameters:

− pruning thresholds (i.e., *α* reference significance levels) for controlling edge formation and the subsequent sparsity of the produced graphs;
− the contributing graph components for likelihood estimates, whether there is a selective focus on edges (*α* = 0), nodes (*β* = 0), or both components;
− aggregation functions (product versus cumulative) and weighting scheme for each node and edge contribution (uniform versus non-uniform);
− normalization of node and edge weights under a min-max approach or dedicated shift factors.

The proposed graph-based predictor can be seen as an ensemble approach where a collective of graph components contribute to the decision.

## 3 Experimental Setting

The validation process of the proposed methodology is composed of multi-omics data derived from TCGA [27]. Three omic platforms – mRNA, miRNA, and protein abundance – were retrieved for five cancer types – Colon Adenocarcinoma (COAD), Kidney Renal Clear Cell Carcinoma (KIRC), Low-Grade Glioma (LGG), Lung Adenocarcinoma (LUAD), and Ovarian Cancer (OV).

### 3.1 Preprocessing

Basic preprocessing steps were applied to all datasets to ensure data quality and consistency. Features with zero variance, missing values, or sparsity exceeding 95% were systematically removed.

Additionally, miRNA data were preprocessed to mitigate cross-mapping biases from impacting the analysis. Cross-mapping occurs when RNA sequences originating from one locus are inadvertently mapped to another [32]. The miRNA molecules with more than 50 cross-mapping events were filtered out to ensure a more accurate representation of the data.

The datasets were split into separate files based on data type and normalization strategy to streamline the analysis. For miRNA data, reads per million mapped reads (RPM) normalization was used, while for mRNA data, transcripts per million (TPM) normalization was applied. Proteomics data were derived from antibody-based high-throughput Reverse-Phase Protein Array (RPPA) technology. Differential Gene Expression (DGE) filtering [33] was applied to the transcriptomic data – mRNA and miRNA – to reduce dimensionality by statistically testing variations in expression across targets, thereby ensuring sensitivity to the inherent stochasticity of the regulatory activity in cancer tissue.

### 3.2 Targets

In this study, we focused on two main outcomes, the *vital status*, a binary variable that indicates the survivability of patients, and the *primary site of the tumor*. These target variables are often imbalanced and, in some cases, were simplified to improve the downstream analysis.

Regarding primary tumor sites, some values were underrepresented due to the low number of associated samples. These were either grouped under broader primary site labels or, when infeasible, discarded. In the LGG dataset, primary sites were grouped into three categories: astrocytoma (anaplastic and not otherwise specified), oligodendroglioma (anaplastic and not otherwise specified), and mixed gliomas. For the COAD dataset, the main categories were adenocarcinoma (NOS) and mucinous adenocarcinoma, excluding rare or ambiguous subtypes with very few samples. Similarly, KIRC data were simplified to two subtypes: clear cell adenocarcinoma (523 samples) and renal cell carcinoma (14 samples), reflecting a severely imbalanced setting. In the LUAD dataset, there are six major subtypes: adenocarcinoma (NOS), adenocarcinoma with mixed subtypes, papillary adenocarcinoma (NOS), acinar cell carcinoma, bronchiolo-alveolar carcinoma (non-mucinous), and mucinous adenocarcinoma. Smaller, ambiguous, or undefined subtypes were excluded to focus on meaningful classifications. Finally, in the OV dataset, the overwhelming majority of cases were classified as serous cystadenocarcinoma with rare variants, resulting in the exclusion of this dataset for the outcome prediction.

### 3.3 Tools and Software

The code used to conduct the experiments in this article is publicly available in the project’s repository^1^. It includes pre-computed graphs derived from the TCGA’s multi-omics data analysis and detailed instructions for running the code, enabling the replication of experiments and further inspection of graph-based predictors.

Data acquisition is performed using the TCGABiolinks package for R, which facilitates the retrieval and preprocessing of datasets from TCGA [34]. Differential Gene Expression analysis is conducted using the edgeR package to identify statistically significant features across outcome classes [33]. The graph generation and prediction processes are entirely original and were implemented in Python, relying on widely adopted libraries. Numpy and Pandas handle numerical tabular data manipulation, while Scipy is employed for statistical computations, including Kolmogorov-Smirnov testing [35, 36, 37]. Scikit-Learn provides essential tools for machine learning workflows, such as the implementations of baseline ML models and evaluation metrics [30].

## 4 Results

The predictive and descriptive capacity of the proposed graph-based structures is now assessed against machine learning baselines using different omics and cancers in TCGA, according to two target variables – the vital status of the patient and the primary tumor tumor site.

The predictive efficacy was assessed under a 5-fold cross-validation schema, with statistical differences tested under a *t*-test. For a conservative view on the performance improvements of the proposed graph-based approaches, the best ML baseline is first identified for each assessed setting (i.e., best predictor for each task, omics layer, and cancer project), and the differences in performance are only assessed against the *best* performing baseline.

### Patient Vital Status Prediction

Table 1 presents the results obtained by the best graph-based configurations for each cancer type and molecular layer. Different parameter combinations are highlighted, such as pruning thresholds and weighting strategies. The best-performing machine learning baseline models are presented in Table 2. All results are averaged across five cross-validation folds. Additionally, values that outperform their ML or graph-based counterparts are highlighted in **bold**, and statistically significant differences are indicated with an asterisk (*).

**Table 1:** Best graph-based configurations for vital status prediction in each cancer project. Hyperparameters considered: i) pruning threshold, ii) graph components used for prediction (nodes, edges, or both), and iii) node/edge weighting scheme (uniform vs. non-uniform).

| Project | Layer | Pruning | Classification | Weighted | Accuracy | F<sub>1</sub> score | Recall | Precision | AUC |
| --- | --- | --- | --- | --- | --- | --- | --- | --- | --- |
| COAD | miRNA | 10<sup>-3</sup> | Both | True | **0.526 ± 0.124** | 0.321 ± 0.052 | 0.511 ± 0.216 | **0.272 ± 0.091** | 0.511 ± 0.054 |
| COAD | mRNA | 10<sup>-2</sup> | Both | False | **0.632 ± 0.106** | 0.378 ± 0.078 | **0.509 ± 0.189** | 0.322 ± 0.072 | 0.639 ± 0.152 |
| COAD | Protein | 10<sup>-4</sup> | Both | True | **0.651 ± 0.087** | **0.345 ± 0.139** | 0.424 ± 0.184 | **0.294 ± 0.116** | 0.568 ± 0.101 |
| KIRC | miRNA | 10<sup>-2</sup> | Both | False | **0.684 ± 0.037\*** | **0.524 ± 0.046** | 0.517 ± 0.048 | **0.533 ± 0.058\*** | **0.688 ± 0.055** |
| KIRC | mRNA | 10<sup>-3</sup> | Both | False | 0.703 ± 0.017 | 0.547 ± 0.029 | **0.548 ± 0.065** | 0.552 ± 0.030 | 0.769 ± 0.029 |
| KIRC | Protein | 10<sup>-4</sup> | Both | True | 0.700 ± 0.025 | 0.575 ± 0.047 | 0.596 ± 0.142 | 0.581 ± 0.076 | 0.742 ± 0.059 |
| LGG | miRNA | 10<sup>-4</sup> | Both | True | **0.789 ± 0.031\*** | **0.514 ± 0.079** | 0.461 ± 0.100 | **0.590 ± 0.060\*** | **0.734 ± 0.023** |
| LGG | mRNA | 10<sup>-4</sup> | Both | True | 0.777 ± 0.058 | 0.531 ± 0.156 | **0.538 ± 0.199** | 0.545 ± 0.128 | **0.768 ± 0.086** |
| LGG | Protein | 10<sup>-4</sup> | Both | True | **0.785 ± 0.034** | 0.444 ± 0.107 | 0.378 ± 0.111 | **0.547 ± 0.105** | **0.704 ± 0.098** |
| LUAD | miRNA | 10<sup>-2</sup> | Both | False | **0.566 ± 0.070** | 0.422 ± 0.045 | 0.447 ± 0.113 | 0.423 ± 0.059 | **0.554 ± 0.103** |
| LUAD | mRNA | 10<sup>-2</sup> | Both | False | 0.634 ± 0.068 | 0.518 ± 0.091 | 0.546 ± 0.098 | 0.495 ± 0.092 | 0.639 ± 0.088 |
| LUAD | Protein | 10<sup>-3</sup> | Both | True | **0.631 ± 0.065** | **0.555 ± 0.060** | **0.574 ± 0.090** | **0.546 ± 0.075** | **0.638 ± 0.053** |
| OV | miRNA | 10<sup>-3</sup> | Edge | True | **0.559 ± 0.050\*** | 0.432 ± 0.057 | 0.463 ± 0.090 | **0.410 ± 0.056** | **0.552 ± 0.059** |
| OV | mRNA | 10<sup>-2</sup> | Both | False | 0.638 ± 0.037 | 0.521 ± 0.054 | 0.527 ± 0.076 | 0.520 ± 0.047 | 0.645 ± 0.074 |
| OV | Protein | 10<sup>-4</sup> | Edge | False | **0.598 ± 0.041** | 0.506 ± 0.110 | 0.544 ± 0.196 | 0.491 ± 0.044 | 0.618 ± 0.042 |

\* statistically verified superiority

**Table 2:**
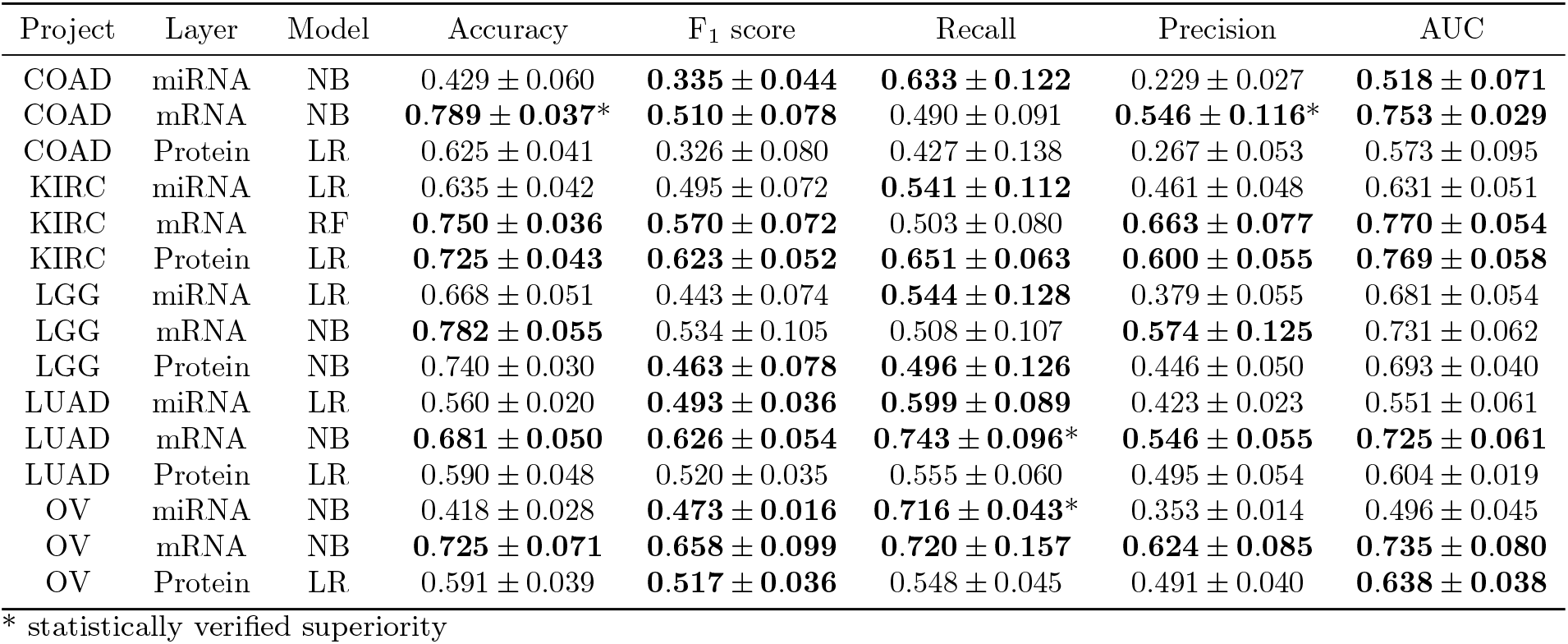
Best ML models for vital status prediction in each cancer project.

| Project | Layer | Model | Accuracy | F<sub>1</sub> score | Recall | Precision | AUC |
| --- | --- | --- | --- | --- | --- | --- | --- |
| COAD | miRNA | NB | 0.429 ± 0.060 | **0.335 ± 0.044** | **0.633 ± 0.122** | 0.229 ± 0.027 | **0.518 ± 0.071** |
| COAD | mRNA | NB | **0.789 ± 0.037\*** | **0.510 ± 0.078** | 0.490 ± 0.091 | **0.546 ± 0.116\*** | **0.753 ± 0.029** |
| COAD | Protein | LR | 0.625 ± 0.041 | 0.326 ± 0.080 | 0.427 ± 0.138 | 0.267 ± 0.053 | 0.573 ± 0.095 |
| KIRC | miRNA | LR | 0.635 ± 0.042 | 0.495 ± 0.072 | **0.541 ± 0.112** | 0.461 ± 0.048 | 0.631 ± 0.051 |
| KIRC | mRNA | RF | **0.750 ± 0.036** | **0.570 ± 0.072** | 0.503 ± 0.080 | **0.663 ± 0.077** | **0.770 ± 0.054** |
| KIRC | Protein | LR | **0.725 ± 0.043** | **0.623 ± 0.052** | **0.651 ± 0.063** | **0.600 ± 0.055** | **0.769 ± 0.058** |
| LGG | miRNA | LR | 0.668 ± 0.051 | 0.443 ± 0.074 | **0.544 ± 0.128** | 0.379 ± 0.055 | 0.681 ± 0.054 |
| LGG | mRNA | NB | **0.782 ± 0.055** | 0.534 ± 0.105 | 0.508 ± 0.107 | **0.574 ± 0.125** | 0.731 ± 0.062 |
| LGG | Protein | NB | 0.740 ± 0.030 | **0.463 ± 0.078** | **0.496 ± 0.126** | 0.446 ± 0.050 | 0.693 ± 0.040 |
| LUAD | miRNA | LR | 0.560 ± 0.020 | **0.493 ± 0.036** | **0.599 ± 0.089** | 0.423 ± 0.023 | 0.551 ± 0.061 |
| LUAD | mRNA | NB | **0.681 ± 0.050** | **0.626 ± 0.054** | **0.743 ± 0.096\*** | **0.546 ± 0.055** | **0.725 ± 0.061** |
| LUAD | Protein | LR | 0.590 ± 0.048 | 0.520 ± 0.035 | 0.555 ± 0.060 | 0.495 ± 0.054 | 0.604 ± 0.019 |
| OV | miRNA | NB | 0.418 ± 0.028 | **0.473 ± 0.016** | **0.716 ± 0.043\*** | 0.353 ± 0.014 | 0.496 ± 0.045 |
| OV | mRNA | NB | **0.725 ± 0.071** | **0.658 ± 0.099** | **0.720 ± 0.157** | **0.624 ± 0.085** | **0.735 ± 0.080** |
| OV | Protein | LR | 0.591 ± 0.039 | **0.517 ± 0.036** | 0.548 ± 0.045 | 0.491 ± 0.040 | **0.638 ± 0.038** |

\* statistically verified superiority

Considering the proposed graph-based representations (Table 1), we observe that the information contained in edges (class-conditional likelihoods of testing ratios) is selected by the hyperparameterization procedure for all predictive settings (cancers and omics layers), confirming the relevance of pairwise class-discriminative relationships in the data. Node information (class-conditional likelihoods of testing features) is disregarded in specific settings. The low pruning thresholds reveal that a comprehensive edge exploration is important to answer the targeted tasks in light of their complexity.

The performance of graph-based approaches is competitive with the best ML baselines. across application contexts, namely the cancer type and the molecular layer used as input. While their overall performance is comparable, there are slight differences to note. In the **KIRC** dataset, the graph-based method demonstrated strong prediction capabilities, achieving the best results (F1, accuracy, AUC) using the mRNA layer. The miRNA-relying variant achieved statistically significant improvements against the best ML baseline, namely on accuracy and precision. The LR model on proteomics data shows competitive predictive performance, although the improvements are not statistically significant. For the **LGG** dataset, the selected comparative example in Figure 2, the proposed graph-based models trained on miRNA data provided the best results, yielding statistically significant differences against the ML baselines in accuracy and precision. Finally, for the remaining datasets, **COAD, LUAD**, and **OV**, we observed competitive behavior between the proposed graph-based approaches and best ML peers, with the choices essentially depending on the selected omics layer and performance metric.

**Figure 2:**
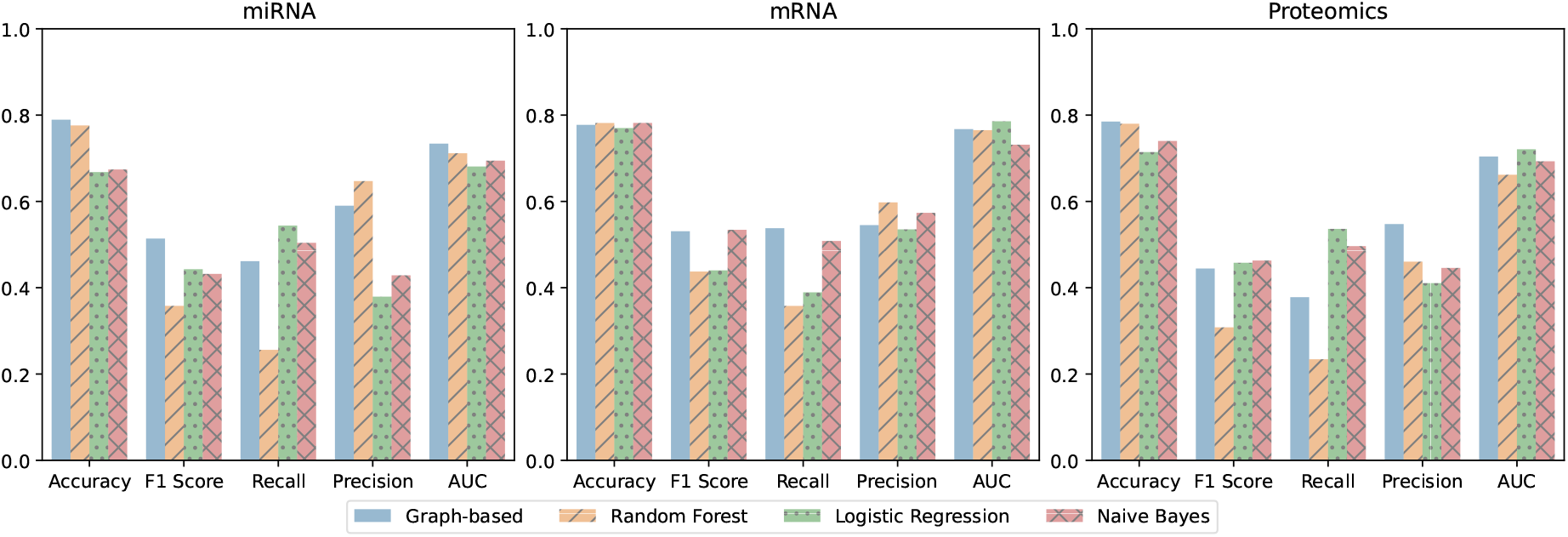
Performance comparison between the best graph-based predictor and the corresponding best ML baseline when predicting the vital status with different omics layers (miRNA, mRNA, proteomics) as input on the Low Grade Glioma (LGG) project.

### Tumor Primary Site Prediction

When tackling primary site prediction, the results further confirm that the predictive performance of graph-based approaches (Table 3) is competitive, superior in diverse settings, against the best ML approaches (Table 4). As before, the performance of the best graph-based configurations is shown in Table 3. Similarly, the annotation of edges with the class-conditional probability density distribution of log-ratios yields relevant predictive information, as highlighted by the selection of edge content by the hyperparameterization. The extent of pruning varies across layers, with higher pruning thresholds being understandably observed for omics with higher dimensionality, particularly transcriptomics.

**Table 3:** Best graph-based configurations for tumor primary site prediction in each cancer project. Hyperparameters considered: i) pruning threshold, ii) graph components used for prediction (nodes, edges, or both), and iii) node/edge weighting scheme (uniform vs. non-uniform).

| Project | Layer | Pruning | Classification | Weighted | Accuracy | F<sub>1</sub> score | Recall | Precision | AUC |
| --- | --- | --- | --- | --- | --- | --- | --- | --- | --- |
| COAD | miRNA | $10^{-3}$ | Both | True | **$0.747 \pm 0.057^*$** | **$0.474 \pm 0.124^*$** | **$0.778 \pm 0.222$** | **$0.342 \pm 0.088^*$** | **$0.790 \pm 0.061^*$** |
| COAD | mRNA | $10^{-2}$ | Both | False | $0.803 \pm 0.047$ | **$0.523 \pm 0.103$** | **$0.756 \pm 0.165$** | $0.402 \pm 0.080$ | **$0.871 \pm 0.086$** |
| COAD | Protein | $10^{-4}$ | node | True | **$0.804 \pm 0.023$** | $0.342 \pm 0.161$ | $0.443 \pm 0.264$ | **$0.287 \pm 0.109$** | $0.710 \pm 0.156$ |
| KIRC | miRNA | $10^{-2}$ | Both | False | **$0.938 \pm 0.036^*$** | **$0.449 \pm 0.156$** | **$0.900 \pm 0.224$** | **$0.332 \pm 0.152$** | **$0.986 \pm 0.014$** |
| KIRC | mRNA | $10^{-2}$ | Both | False | $0.968 \pm 0.026$ | $0.554 \pm 0.384$ | **$0.800 \pm 0.447$** | $0.463 \pm 0.385$ | **$0.957 \pm 0.096$** |
| KIRC | Protein | $10^{-4}$ | Both | True | $0.955 \pm 0.021$ | $0.457 \pm 0.096$ | $1.000 \pm 0.000$ | $0.300 \pm 0.075$ | $0.974 \pm 0.017$ |
| LGG | miRNA | $10^{-4}$ | Both | False | $0.580 \pm 0.028$ | **$0.546 \pm 0.033^*$** | **$0.549 \pm 0.029$** | **$0.547 \pm 0.036$** | **$0.712 \pm 0.034$** |
| LGG | mRNA | $10^{-4}$ | Both | True | **$0.607 \pm 0.050$** | **$0.553 \pm 0.057$** | **$0.564 \pm 0.055$** | **$0.572 \pm 0.072$** | $0.701 \pm 0.043$ |
| LGG | Protein | $10^{-4}$ | Edge | True | $0.528 \pm 0.055$ | **$0.528 \pm 0.054$** | **$0.527 \pm 0.059$** | **$0.565 \pm 0.053$** | $0.694 \pm 0.028$ |
| LUAD | miRNA | $10^{-2}$ | Both | True | $0.168 \pm 0.017$ | $0.174 \pm 0.041$ | **$0.373 \pm 0.067$** | **$0.244 \pm 0.050$** | **$0.636 \pm 0.039$** |
| LUAD | mRNA | $10^{-2}$ | Both | True | **$0.273 \pm 0.051$** | **$0.192 \pm 0.034$** | **$0.264 \pm 0.071$** | **$0.277 \pm 0.063$** | **$0.613 \pm 0.040$** |
| LUAD | Protein | $10^{-4}$ | Edge | True | $0.196 \pm 0.054$ | $0.091 \pm 0.041$ | $0.124 \pm 0.077$ | $0.149 \pm 0.039$ | $0.477 \pm 0.060$ |

\* statistically verified superiority

**Table 4:** Best ML models for tumor primary site prediction in each cancer project.

| Project | Layer | Model | Accuracy | F<sub>1</sub> score | Recall | Precision | AUC |
| --- | --- | --- | --- | --- | --- | --- | --- |
| COAD | miRNA | LR | 0.609 ± 0.038 | 0.290 ± 0.022 | 0.549 ± 0.095 | 0.199 ± 0.010 | 0.626 ± 0.036 |
| COAD | mRNA | NB | **0.855 ± 0.022\*** | 0.519 ± 0.058 | 0.538 ± 0.084 | **0.513 ± 0.094\*** | 0.850 ± 0.039 |
| COAD | protein | LR | 0.749 ± 0.061 | **0.380 ± 0.123** | **0.608 ± 0.196** | 0.279 ± 0.095 | **0.762 ± 0.059** |
| KIRC | miRNA | NB | 0.879 ± 0.038 | 0.267 ± 0.089 | 0.833 ± 0.211 | 0.163 ± 0.062 | 0.830 ± 0.152 |
| KIRC | mRNA | NB | **0.979 ± 0.011** | **0.573 ± 0.177** | 0.567 ± 0.249 | **0.667 ± 0.279** | 0.803 ± 0.158 |
| KIRC | protein | NB | **0.973 ± 0.020** | **0.629 ± 0.232** | 1.000 ± 0.000 | **0.507 ± 0.288** | **0.986 ± 0.010** |
| LGG | miRNA | RF | **0.608 ± 0.019** | 0.481 ± 0.020 | 0.542 ± 0.017 | 0.468 ± 0.064 | 0.698 ± 0.041 |
| LGG | mRNA | NB | 0.556 ± 0.058 | 0.545 ± 0.059 | 0.544 ± 0.060 | 0.565 ± 0.054 | **0.736 ± 0.039** |
| LGG | protein | LR | **0.549 ± 0.037** | 0.511 ± 0.039 | 0.519 ± 0.035 | 0.515 ± 0.039 | **0.724 ± 0.018** |
| LUAD | miRNA | NB | **0.227 ± 0.038\*** | 0.179 ± 0.040 | 0.335 ± 0.087 | 0.227 ± 0.034 | 0.604 ± 0.066 |
| LUAD | mRNA | NB | 0.277 ± 0.038 | 0.185 ± 0.053 | 0.253 ± 0.083 | 0.236 ± 0.045 | 0.605 ± 0.066 |
| LUAD | protein | NB | **0.453 ± 0.050\*** | **0.208 ± 0.056\*** | **0.226 ± 0.068** | **0.212 ± 0.054** | **0.585 ± 0.083** |

\* statistically verified superiority

In the **COAD** dataset, the graph-based approach demonstrates strong performance, particularly when using mRNA data. When applied to miRNA data, it is statistically superior across all metrics, except for recall, which, on average, remains above the machine learning baseline. For **KIRC**, Naive Bayes and graph-based approaches achieve the best overall results using mRNA data. Both methods yield particularly high accuracy and AUC when compared to other experiments. The only statistically significant difference in performance is observed with the graph-based approach using miRNA data, which demonstrates superior accuracy compared to the best ML baseline. For **LGG**, the selected comparative example in Figure 3, the performance difference between the two approaches is minimal. Therefore, defining the best predictor is challenging. While the best ML baselines may outperform in accuracy, they are surpassed in F1-score, with graph-based approaches yielding statistically significantly higher F1-score using miRNA data. In **LUAD**, performance remains comparable, and proteomics-based models achieve the highest accuracy and F1-score values.

**Figure 3:**
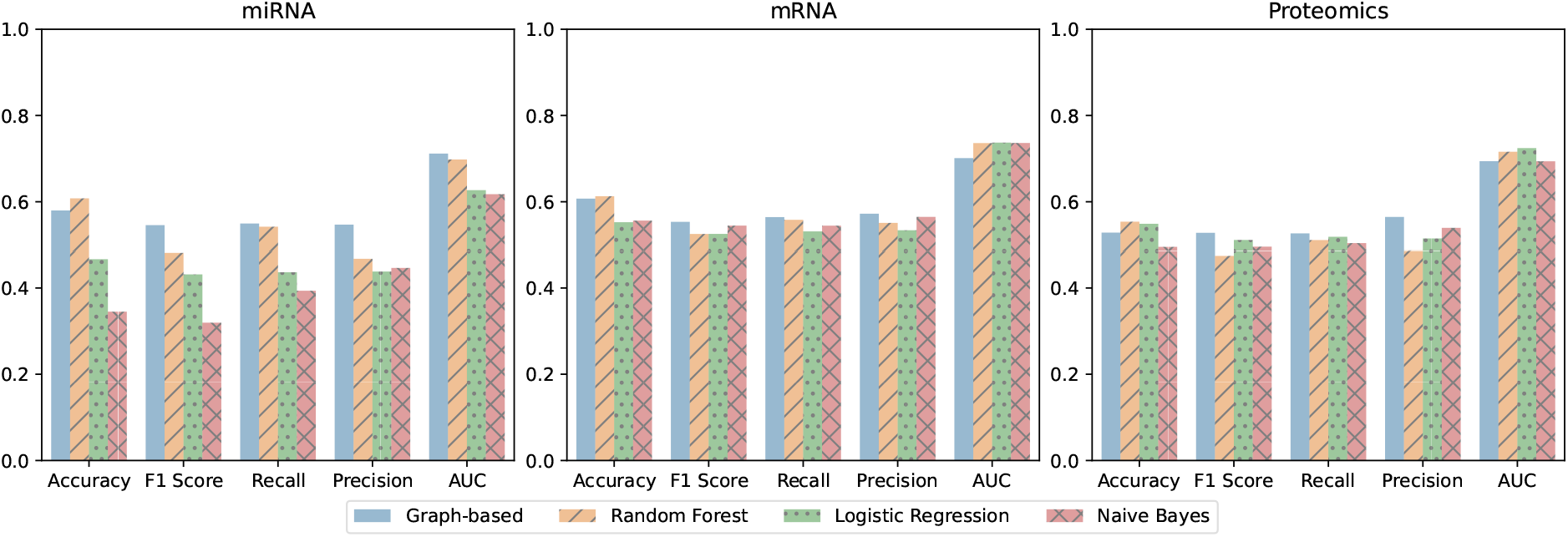
Performance comparison between the best graph-based predictor and the corresponding best ML baseline when predicting the primary tumor site with different omics layers (miRNA, mRNA, proteomics) as input on the Low Grade Glioma (LGG) project.

Complementary results are presented in the supplementary materials. Section S1 extends the graphical analysis of the best predictors towards other cancer projects, section S2 performs a sensitivity analysis of how performance varies with hyperparameterization options, and section S3 acquires insights on the weight distributions of nodes and edges from the produced graphs.

## 5 Discussion

Once an appropriate graph representation is achieved from the available data, it can then be used to extract actionable information for knowledge discovery. This is valid for unsupervised contexts, as well as supervised contexts, where the inspection of original and sample-specific graphs can be used to produce global and local explanations.

In the following subsections, we delve into graph exploration, the effects of parameterizations, and topology analysis (e.g., degree, cliques), as ways of acquiring knowledge. As the detailed analysis of all the datasets and configurations would be far too extensive, the LGG cancer project and the proteomic layer are selected as a guiding case study.

### Pruning

An important step in the proposed methodology is filtering the final graph by retaining only edges that exhibit statistically significant differences between class-conditional distributions. This process enhances interpretability and ensures that preserved edges show guarantees of discriminative power. Additionally, the filtering step controls graph density through the selection of a *p*-value threshold. Lower thresholds result in sparser graphs, preserving only the most distinctive relationships between classes.

Pruning also plays a critical role in predictive performance. Some of the best results are achieved with more aggressive pruning as seen in Tables 1 and 3. The substantial reduction in edges helps eliminate noise while preserving the most relevant structural information. Table 5 summarizes the impact of different *p*-value thresholds on the graph structure, illustrating how increasing pruning levels affect node and edge retention for the LGG dataset using proteomics data.

**Table 5:** Graph attributes according to statistical significance pruning – an example from the LGG project using proteomics data.

| $p$-value | Nodes | % Node Reduction | Edges | % Edge Reduction |
| --- | --- | --- | --- | --- |
| 0.01 | 456 | 0 | 31896 | 69.25 |
| 0.001 | 454 | 0.44 | 16914 | 83.70 |
| 0.0001 | 450 | 1.32 | 8109 | 92.18 |

### Parameterization Effects

According to Tables 1 and 3, predictive strategies that incorporate both node and edge information generally outperform those relying on a single structure, as they leverage a broader range of information. In a few cases, peaked when relying solely on edges. Despite these exceptions, the results remain comparable to approaches that use a combination of both structures, where edges represent the majority of information due to a higher amount of connections relative to the number of nodes in the graphs, as confirmed in Table 5.

Regarding **scaling**, the vast majority of configurations seem to perform better when some type of scaling is applied to both node and edge weights. Similarly, **weighting** generally yields improvements in predictive performance due to a greater capacity to differentiate the contributions across nodes and edges in accordance with their discriminative capacity as given by the KS statistic.

Finally, when considering the input **layers** as a controllable hyperparameter, the best results, especially considering the F_1_ score, are often achieved with mRNA data. This can be verified in both graph-based and ML approaches, as seen in Tables 1 to 4. The only exceptions reside in the LUAD and KIRC projects, where the ML models achieved a higher performance using proteomics data. Interestingly, when considering miRNA data alone, graph-based approaches are able to generally sustain their levels of performance, while the baseline ML models show statistically significant deterioration in performance using data from this specific layer as input.

### Degree Analysis

Figure 4 depicts the degree distribution of the graph inferred from the proteomics layer of the LGG project. This topological analysis suggests that, across classes, most proteins have distinct relationships with only a small number of other proteins, with only a few hub proteins showing altered interactions with more than 300 others.

**Figure 4:**
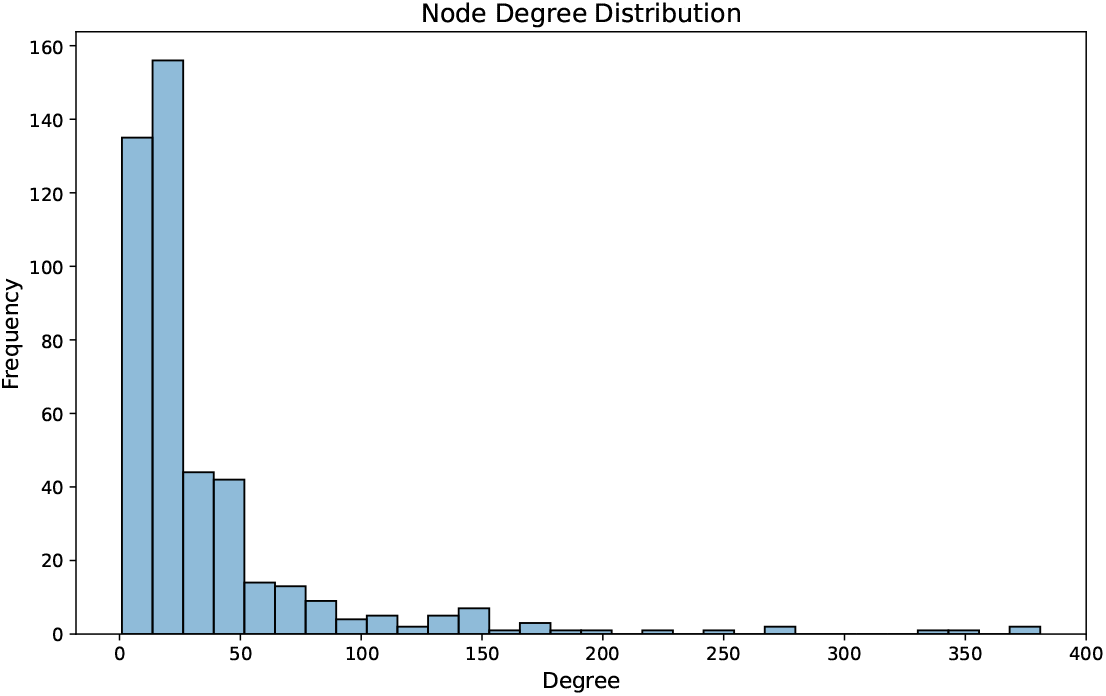
Degree distribution for the nodes in the LGG proteomics graph (0.01 *p*-value pruning).

High-degree nodes may represent biological entities that are involved in numerous significant relationships, making their analysis particularly informative. By filtering nodes with a degree of 200 or higher, a list of 8 proteins is obtained, which are coded by the following genes: BRD4, PPIF, GLUD1, IGFBP2, MMP14, EIF2AK3, PRKCA, and WEE1. A subsequent gene enrichment analysis on these genes reveals their relevance to various processes associated with the Glioma disease, considered in this specific example. Some of the key findings are summarized in Table 6.

**Table 6:** Enrichment terms associated with the gene set: BRD4, PPIF, GLUD1, IGFBP2, MMP14, EIF2AK3, PRKCA, and WEE1.

| Category of Enrichment | Gene Set Library | Enrichment Term |
| --- | --- | --- |
| Pathways | BioPlanet 2019 | MAPK cascade role in angiogenesis |
| Pathways | Elsevier Pathway Collection | Proteins Involved in Glioma |
| Ontologies | GO Biological Process 2023 | Regulation Of Intrinsic Apoptotic Signaling Pathway |
| Diseases/Drugs | DisGeNET | Glioma |

To further assess the relevance of this gene set, an additional statistical analysis was conducted to assess how well the encoded proteins discriminate between the different classes. For each of the proteins associated with the genes selected earlier, a *t*-test is performed to compare their class-conditioned statistical distributions. The results, including the t-statistic, *p*-value, and ranking by *p*-value, are presented in Table 7. These results indicate that proteins associated with high-degree nodes in the graph are also highly discriminative between classes. This validates the methodology’s effectiveness in capturing important, class-separating features.

**Table 7:** Statistical assessment of the class-conditional distributions of a selected set of hub proteins, including their name, *t*-test score, *p*-value, and overall ranking based on statistical significance.

| Gene | $t$-statistic | $p$-value | Ranking |
| --- | --- | --- | --- |
| BRD4 | -5.807 | 1.25e-08 | 4 |
| PPIF | 5.582 | 4.24e-08 | 8 |
| GLUD1 | 5.592 | 4.03e-08 | 7 |
| IGFBP2 | -5.916 | 6.79e-09 | 3 |
| MMP14 | -5.538 | 5.36e-08 | 10 |
| EIF2AK3 | -6.836 | 2.84e-11 | 2 |
| PRKCA | 4.846 | 1.77e-06 | 17 |
| WEE1 | -7.394 | 7.63e-13 | 1 |

### Clique Analysis

The analysis of cliques, subsets of nodes where every node is connected to every other node, is further undertaken on the proposed graph structures for knowledge augmentation. In the context of our case study, a clique represents a group of proteins whose relationships with every other protein of the group show statistically significant capacity to discriminate between LGG classes. This can help identify sets of proteins that form putative functional modules closely related to patient survivability.

The graph under analysis has a maximum clique size of 5, with 424 cliques of this size. These cliques can be further studied through enrichment analysis, as previously done, to gain a better understanding of their functional implications. However, due to the large number of cliques and their relatively small sizes, a detailed analysis of each individual clique becomes challenging, and meaningful results are limited. This limitation arises from the strict definition of a clique, where every node must be connected to every other node. Therefore, relaxations to this task are desirable to identify strongly connected nodes that are not necessarily fully connected.

To address this challenge, the concept of a *k*-core is introduced, which refers to a subset of nodes where each node is connected to at least *k* others within the same subset. Figure 5 illustrates the relationship between the value of *k* for a *k*-core and the number of nodes it contains. From the figure, we can then see that the maximum *k*-core in the graph has *k* = 35. This subgraph can be interpreted as containing the principal nodes and edges in the graph, i.e., those showing statistical guarantees of discriminating outcomes of interest.

**Figure 5:**
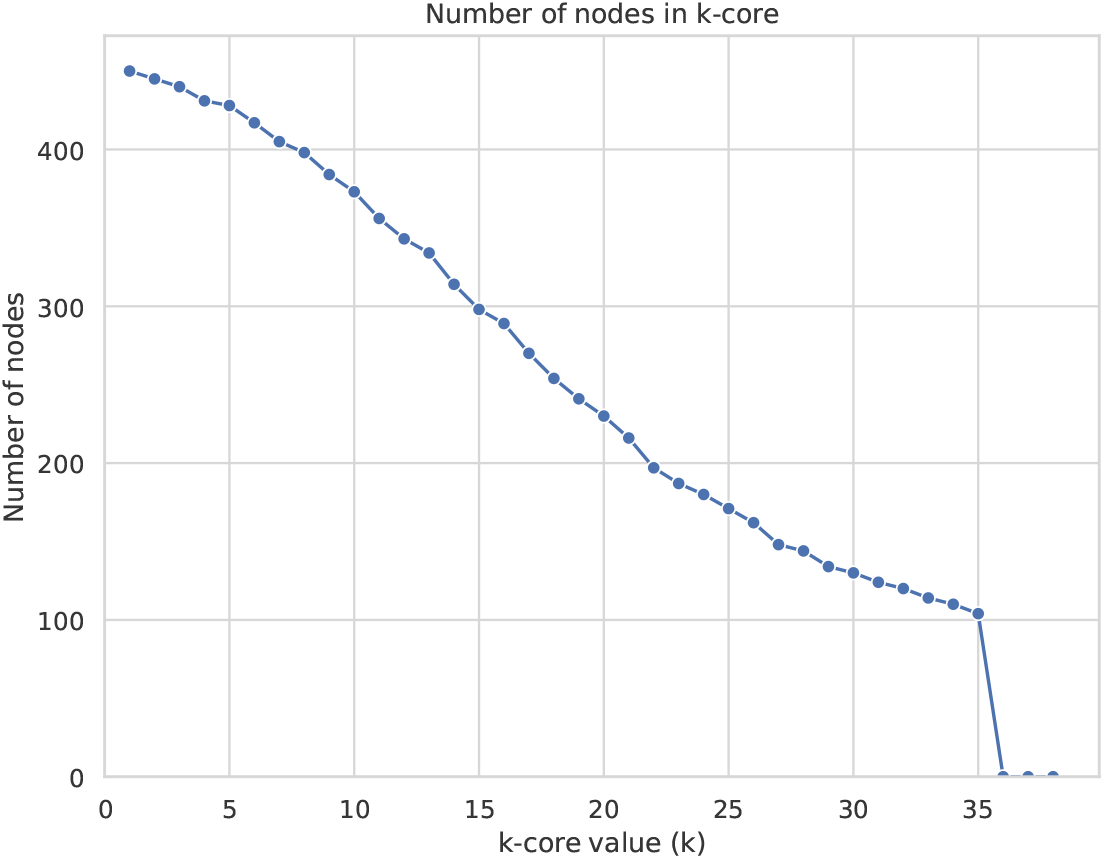
Relationship between the *k* value for a *k*-core and the number of nodes it contains.

## 6 Conclusion

This work proposes an augmented graph-based representation of multi-omics data, along with novel predictive strategies. Our approach focuses on capturing the rich and stochastic nature of relationships between biological entities by encoding class-conditional probability functions in a graph structure.

The proposed methodology shows solid descriptive and predictive learning capacities in multiple oncological contexts. Considering descriptive stances, relationships between entities exhibiting statistically significant alterations across the studied phenotypes are highlighted, contributing to a deeper understanding of the diseases. The analysis of the revised degree distribution scores proved effective in identifying proteins that are both discriminative of the studied phenotypes and, when subjected to enrichment analysis, putatively involved in the development of the target conditions. Complementarily, the analysis of the largest cliques and k-cores within the proposed graph representations allows the identification of multi-wise molecular associations that consistently exhibit changes across phenotypes, suggesting their role as functional modules.

The graphs are further leveraged for predictive tasks, with a focus on classification, by performing kernel density estimation against class-conditional distributions. The utility of this approach is demonstrated across a wide range of predictive TCGA challenges, including predicting patients’ vital status and primary tumor sites across multiple cancer types and various omics layers. The results confirm competitive, at times superior, performance against machine learning models, demonstrating an enhanced ability to handle imbalanced data and offering statistical frames and notable interpretability.

From here, several ramifying paths are highlighted for future work, including the exploration of alternative statistical tests to summarize graph information; complementary contrastive functions on the edges, such as absolute differences or alternative ratios; or additional node/edge filtering strategies that may contribute to more efficient data processing. Future opportunities may also include extensions to accommodate the interplay between omics layers, which will pose new challenges due to increased computational complexity arising from more holistic perspectives. Moreover, the methodology can be extended to handle continuous outcomes by replacing class-conditional distributions with bivariate distributions from paired input-output variables for regression approaches.

## Ethics Statement

This study utilized publicly available datasets. Ethical approval was not required.

## Acknowledgements

The results shown here are in whole or part based upon data generated by the TCGA Research Network: https://www.cancer.gov/tcga.

## Funding

This work was supported by the FCT PhD grant to DG (2022.12633.BD); financed by national funds from FCT - Fundação para a Ciência e a Tecnologia, I.P., in the scope of the project UIDP/04378/2020 (DOI: 10.54499/UIDP/04378/2020) and UIDB/04378/2020 (DOI: 10.54499/UIDB/04378/2020) of the Research Unit on Applied Molecular Biosciences – UCIBIO and the project LA/P/0140/2020 (DOI: 10.54499/LA/P/0140/2020) of the Associate Laboratory Institute for Health and Bioeconomy - i4HB, and INESC-ID plurianual (UIDB/50021/2020); the projects FRAIL (2024.07266.IACDC) and LAIfeBlood+ (2024.07475.IACDC). The authors also wish to acknowledge the European Union’s Horizon BioLaMer project under grant agreement number [101099487]. Any opinions, findings, conclusions, or recommendations expressed in this material are those of the authors and do not necessarily reflect the views of the funding bodies.

## Declaration of Competing Interest

The authors declare that they have no known competing financial interests or personal relationships that could influence the work reported in this paper.

## CRediT Authorship Contribution Statement

**Daniel M. Gonçalves**: Conceptualization, Methodology, Software, Validation, Writing - Original Draft, Writing - Review & Editing; **André Patricio**: Conceptualization, Methodology, Software, Data Curation, Writing - Original Draft; **Rafael S. Costa**: Conceptualization, Methodology, Validation, Writing - Review & Editing, Supervision; **Rui Henriques**: Conceptualization, Methodology, Validation, Writing - Review & Editing, Supervision.

## Supplementary Materials

### S1 Best Predictor Models

**Figure S1:**
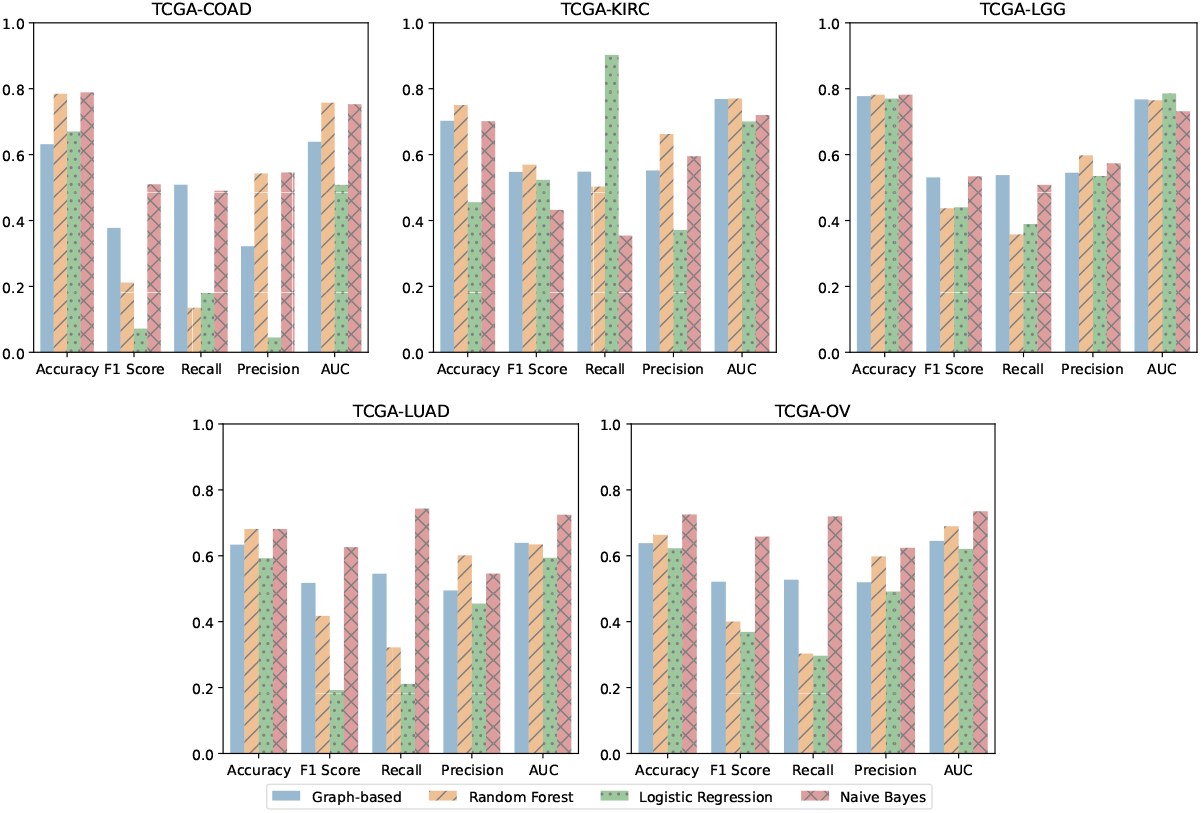
Best graph-based predictor and corresponding ML baselines using protein data to predict the vital status across cancer datasets.

**Figure S2:**
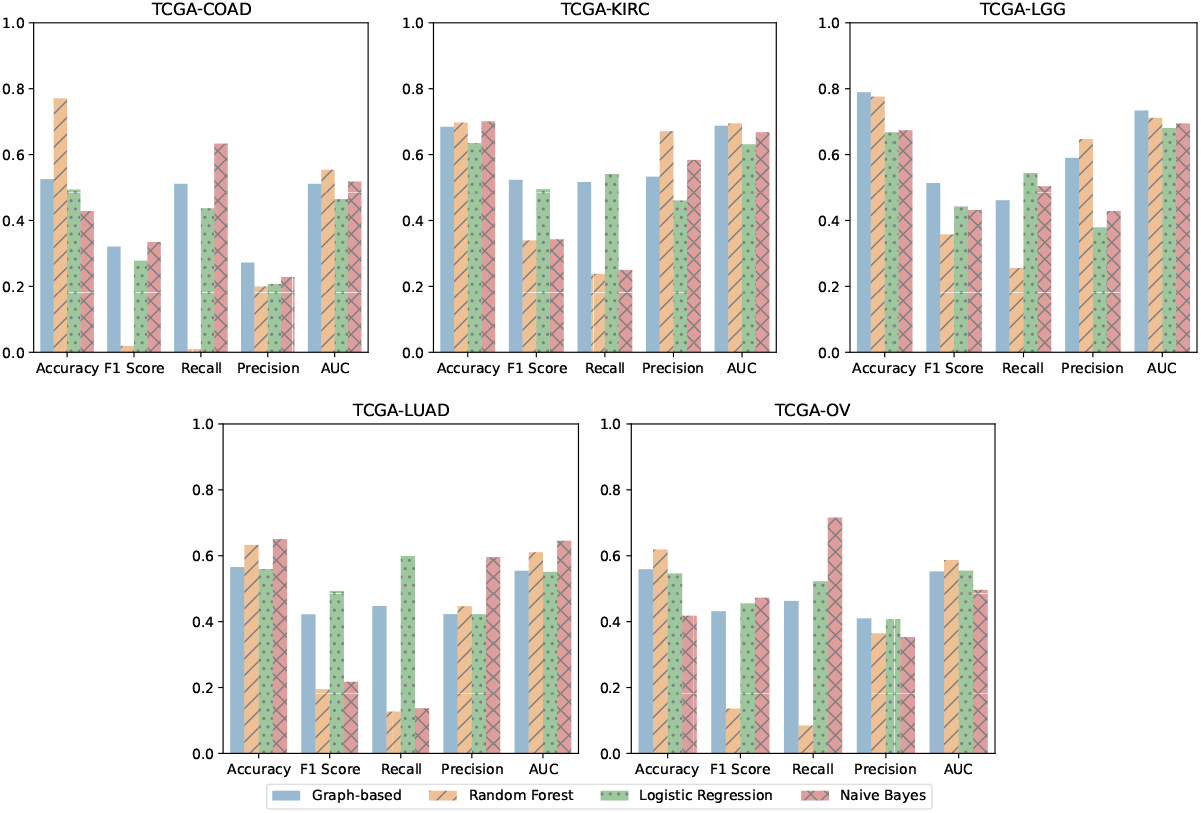
Best graph-based predictor and corresponding ML baselines using protein data to predict the vital status across cancer datasets.

**Figure S3:**
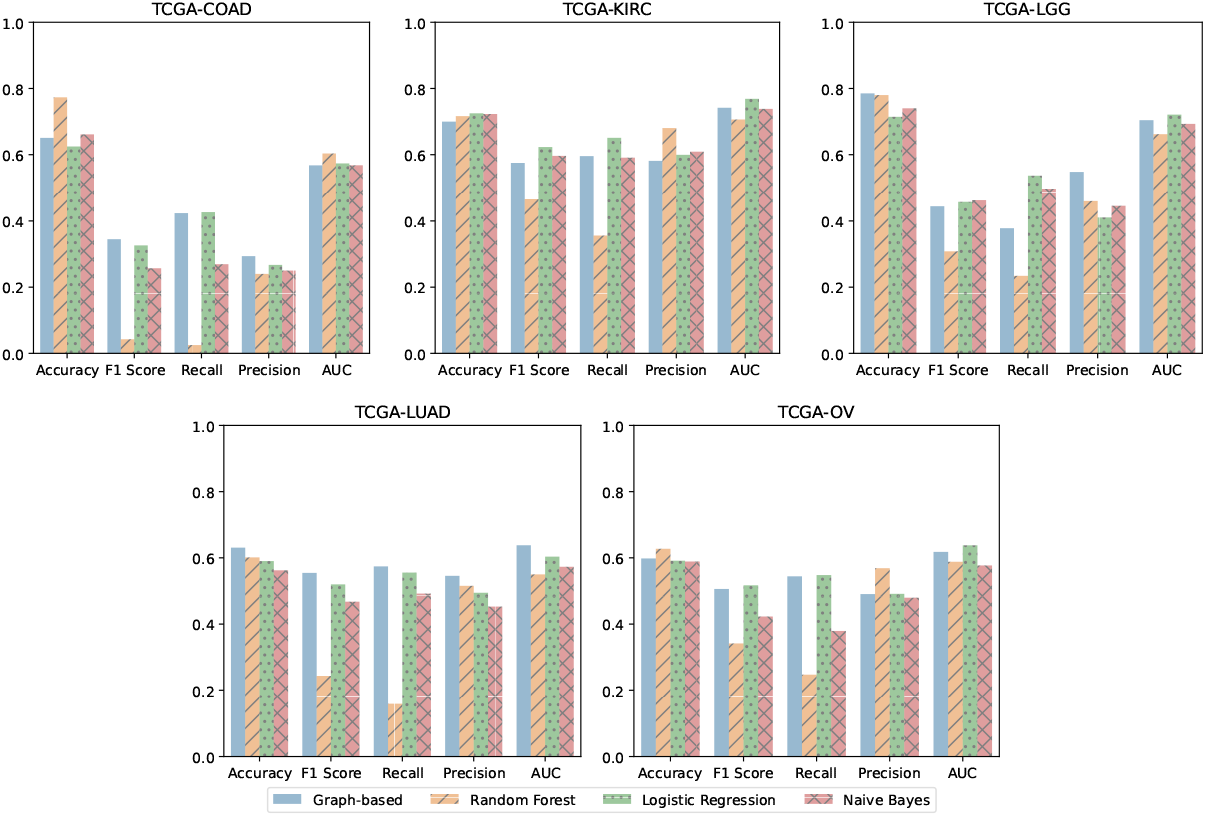
Best graph-based predictor and corresponding ML baselines using protein data to predict the vital status across cancer datasets.

**Figure S4:**
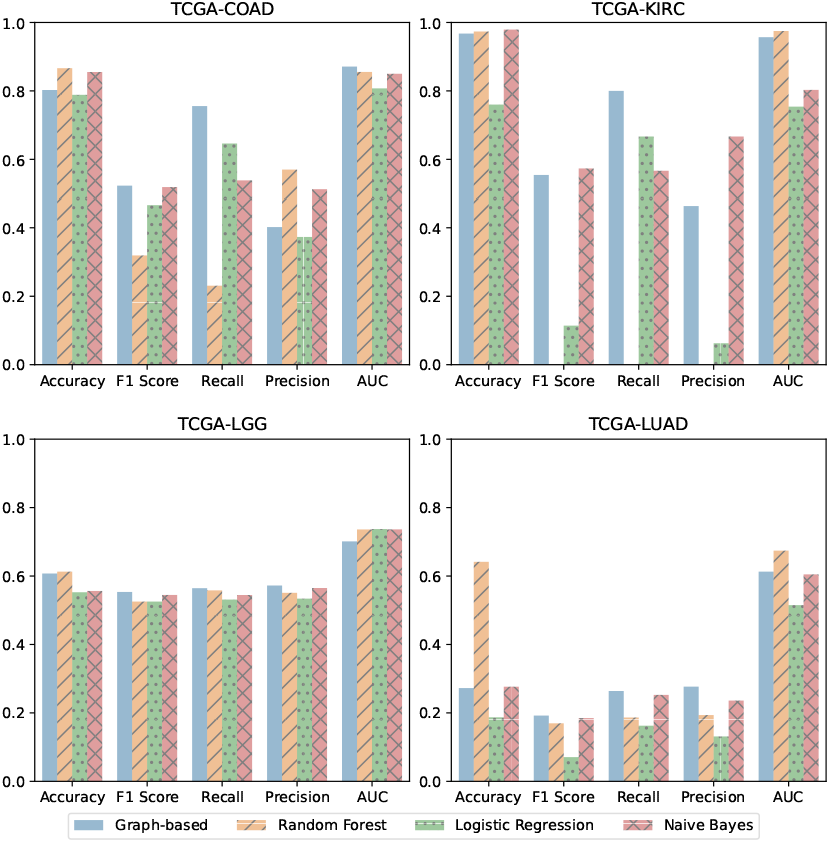
Best graph-based predictor and corresponding ML baselines using mRNA data to predict the primary tumor site across cancer datasets.

**Figure S5:**
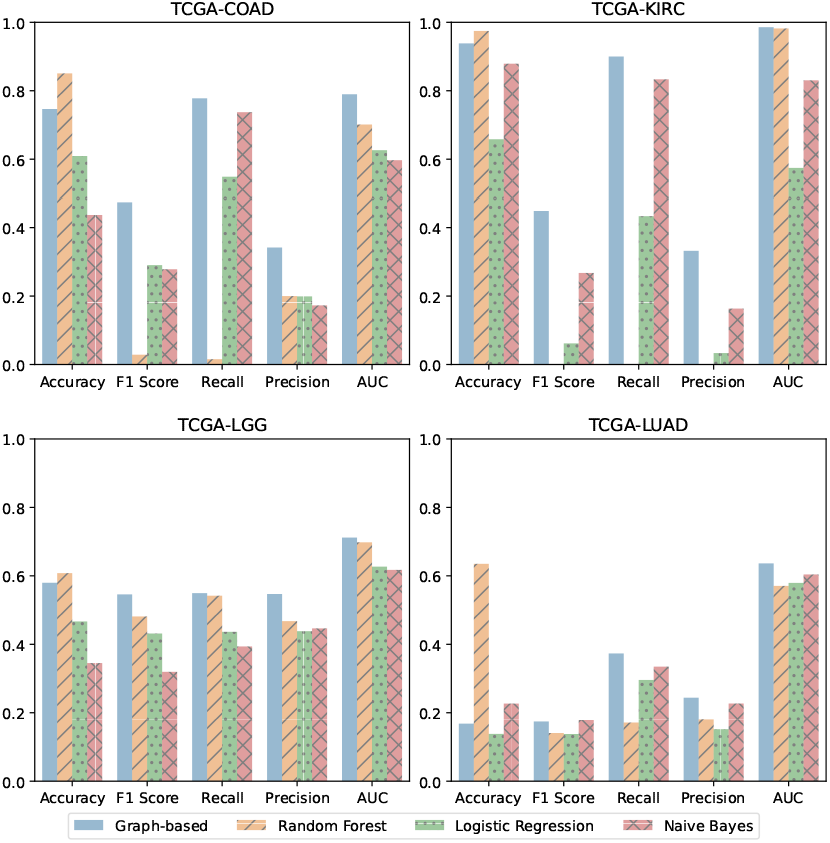
Best graph-based predictor and corresponding ML baselines using miRNA data to predict the primary tumor site across cancer datasets.

**Figure S6:**
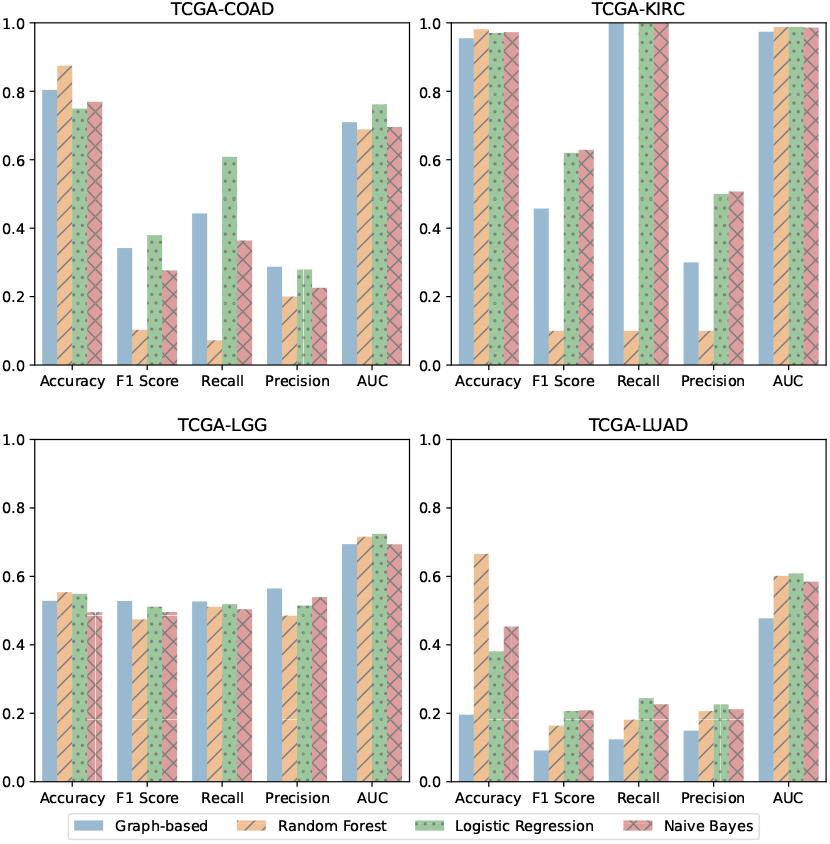
Best graph-based predictor and corresponding ML baselines using protein data to predict the primary tumor site across cancer datasets.

### S2 Parameter Variation Analysis

**Figure S7:**
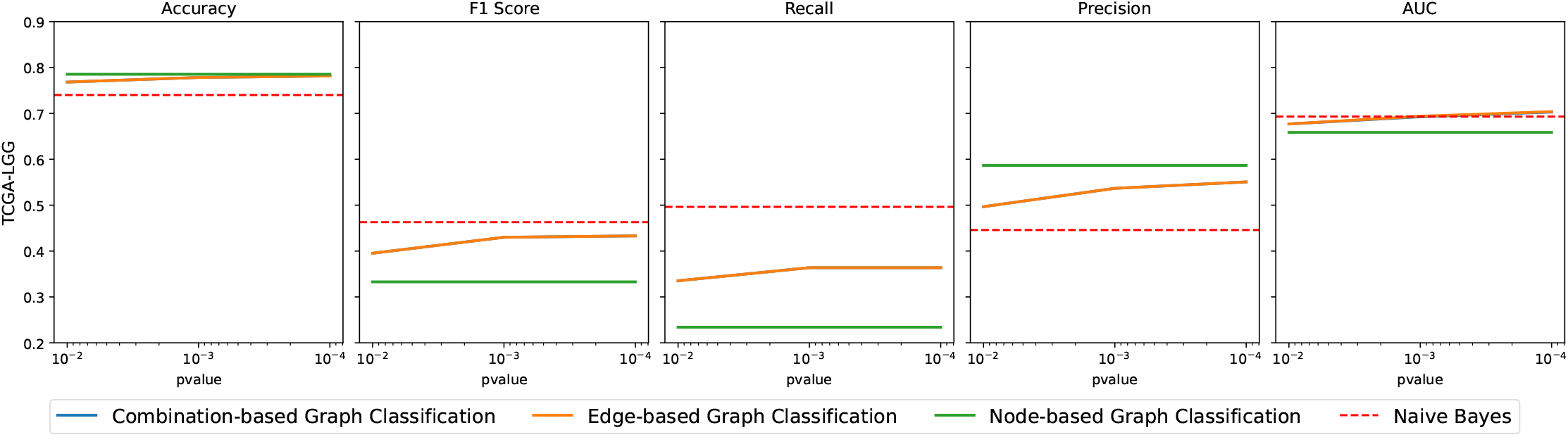
Performance variance of different classification strategies along different pruning thresholds in LGG using proteomics data to predict the vital status.

**Figure S8:**
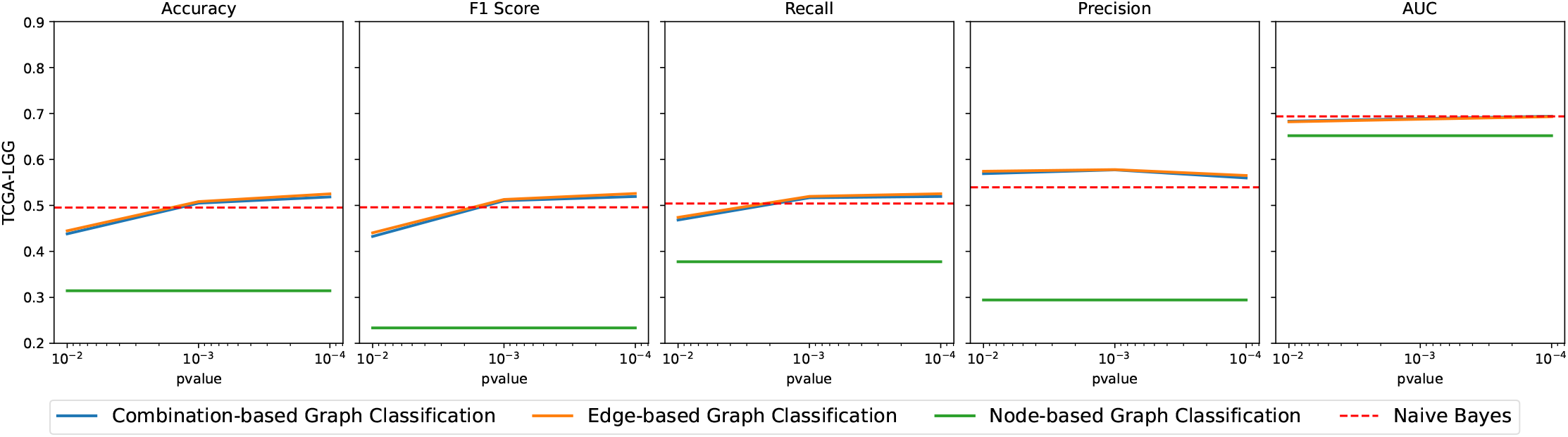
Performance variance of different classification strategies along different pruning thresholds in LGG using proteomics data to predict the primary tumor.

**Figure S9:**
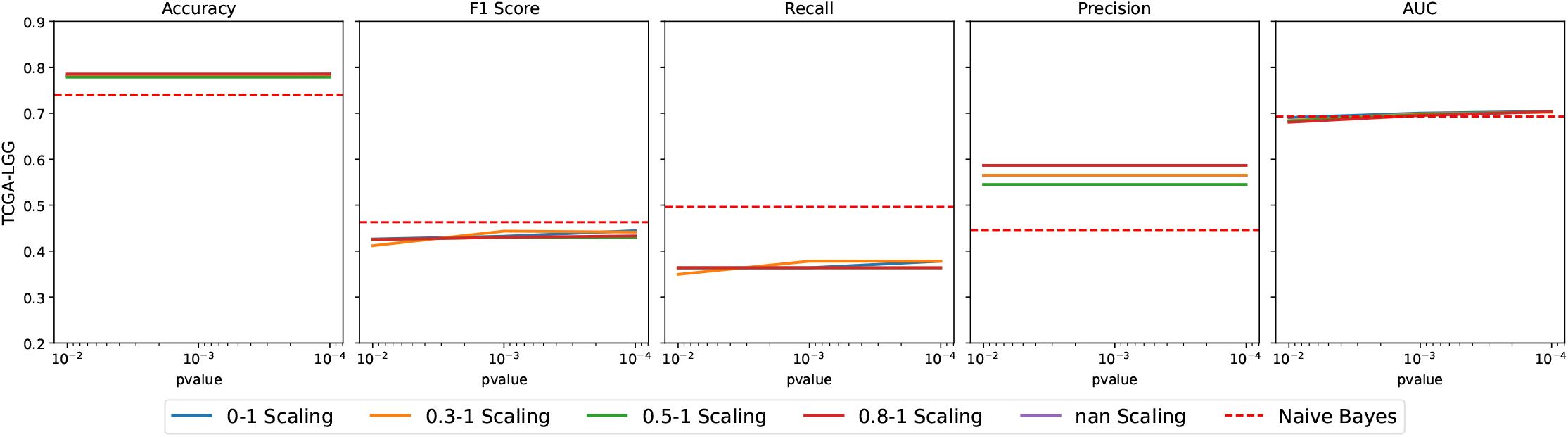
Performance variance of different scaling intervals along different pruning thresholds in LGG using proteomics data to predict the vital status.

**Figure S10:**
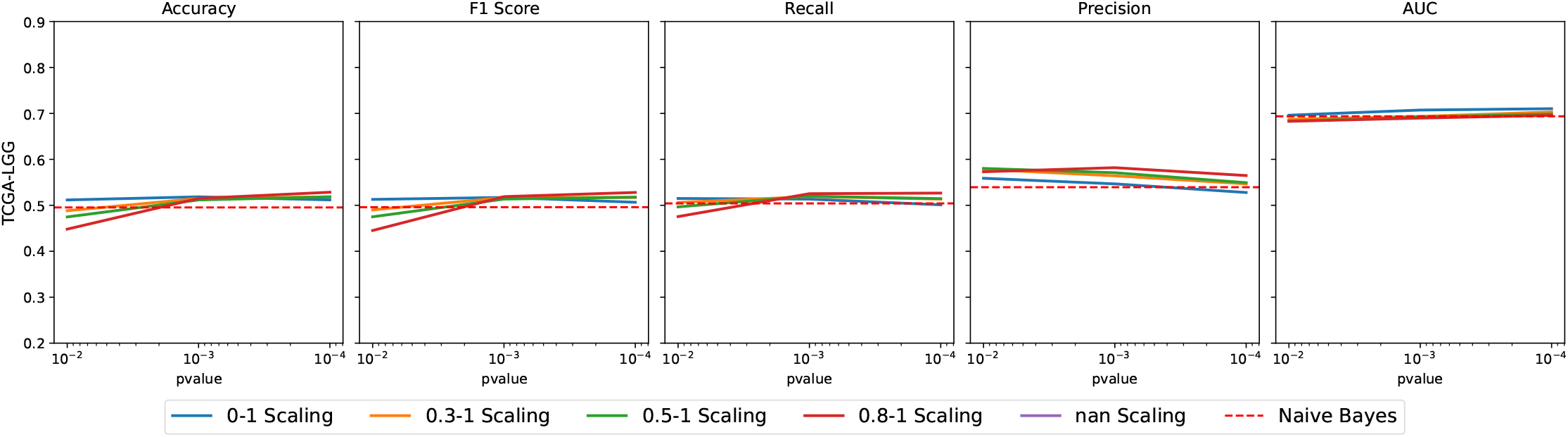
Performance variance of different scaling intervals along different pruning thresholds in LGG using proteomics data to predict the primary tumor.

### S3 Graph Weight Distributions

**Figure S11:**
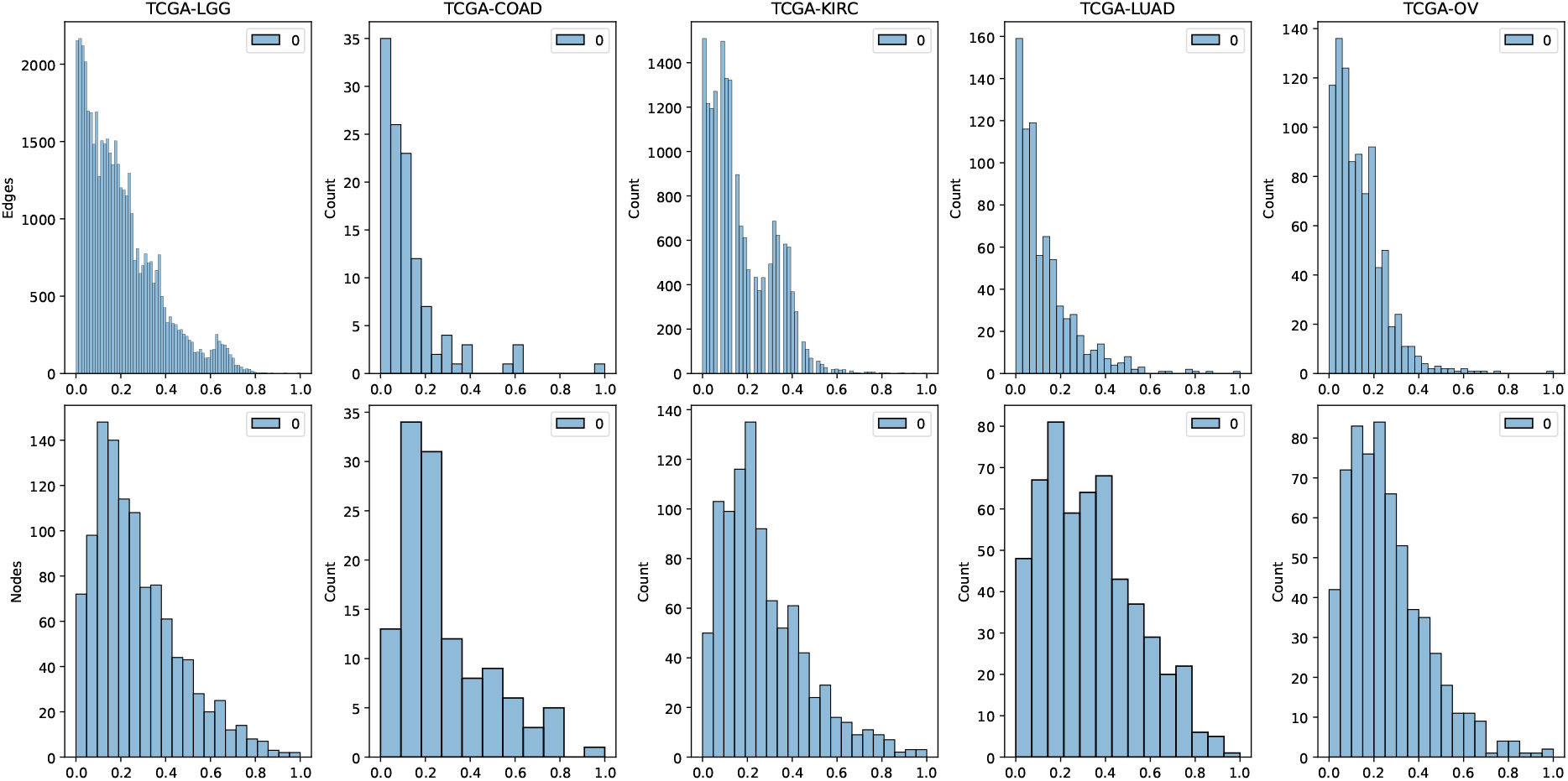
Weight value distribution histograms per cancer project, showing edge and node perspectives using miRNA data.

**Figure S12:**
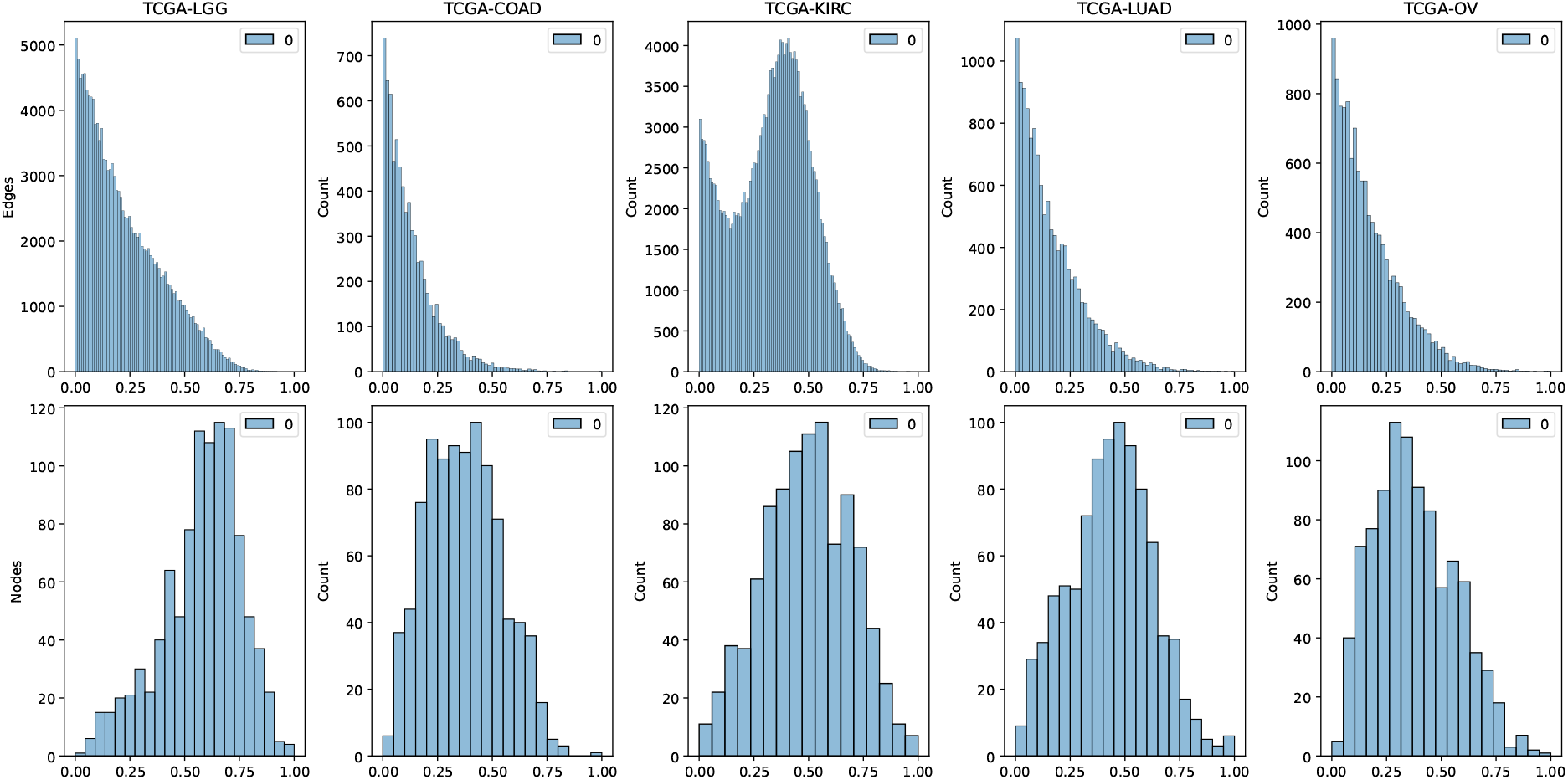
Weight value distribution histograms per cancer project, showing edge and node perspectives using mRNA data.

**Figure S13:**
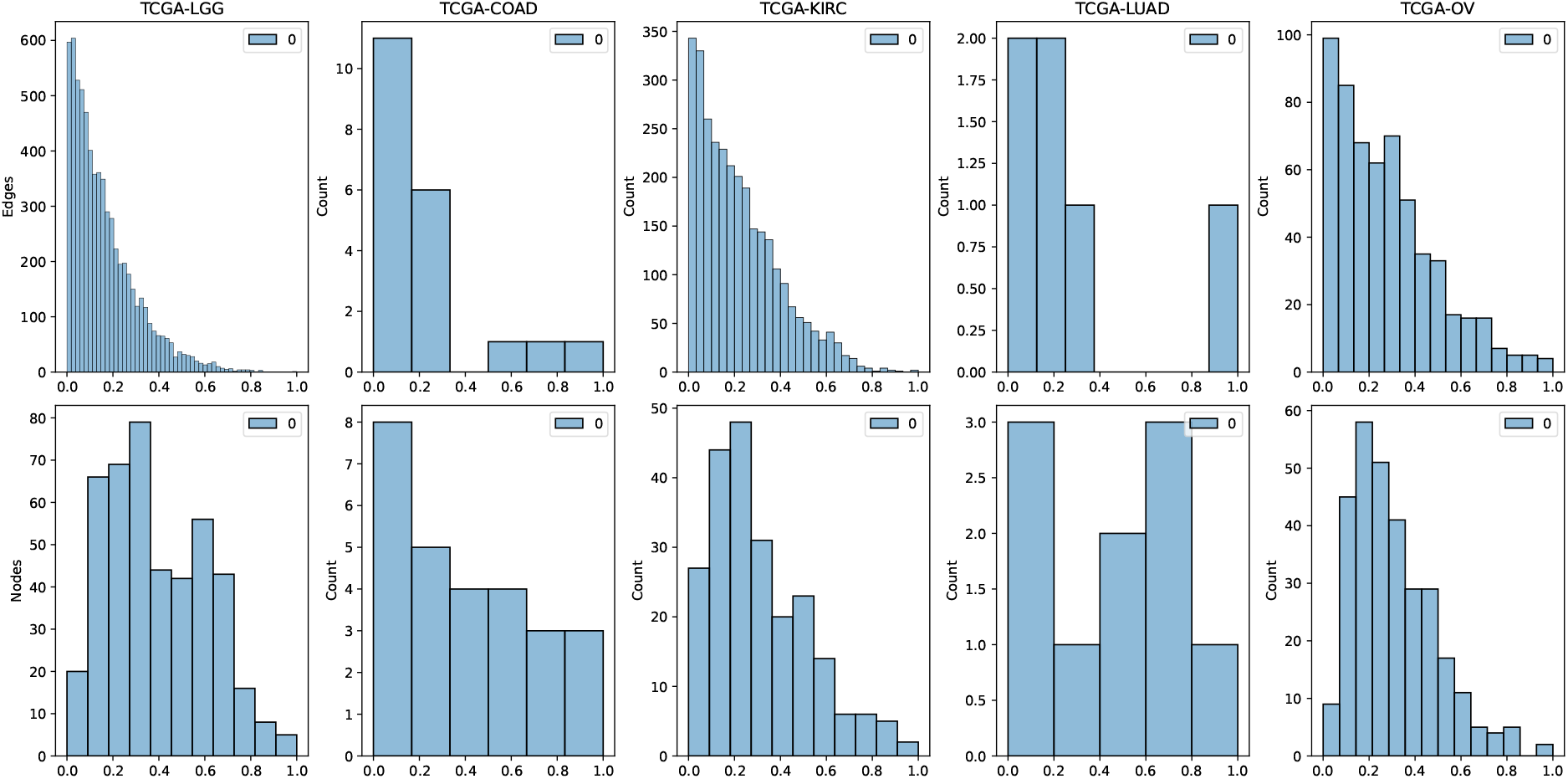
Weight value distribution histograms per cancer project, showing edge and node perspectives using proteomics data.

## Footnotes

1 github.com/dmgoncal/statgraph-omics

## Notes

### Competing Interest Statement

The authors have declared no competing interest.

## References

[1] Douglas Hanahan and Robert A Weinberg. Hallmarks of cancer: the next generation. cell, 144(5):646–674, 2011.

[2] Rui Chen and Michael Snyder. Promise of personalized omics to precision medicine. Wiley Interdisciplinary Reviews: Systems Biology and Medicine, 5(1):73–82, 2013.

[3] Albert-Lászl Barabasi, Natali Gulbahce, and Joseph Loscalzo. Network medicine: a network-based approach to human disease. Nature reviews genetics, 12(1):56–68, 2011.

[4] Lenore Cowen, Trey Ideker, Benjamin J Raphael, and Roded Sharan. Network propagation: a universal amplifier of genetic associations. Nature Reviews Genetics, 18(9):551–562, 2017.

[5] Leroy Hood and Lee Rowen. The human genome project: big science transforms biology and medicine. Genome medicine, 5:1–8, 2013.

[6] Mark Reimers and Vincent J Carey. [8] bioconductor: an open source framework for bioinformatics and computational biology. Methods in enzymology, 411:119–134, 2006.

[7] Karsten M Borgwardt, Cheng Soon Ong, Stefan Schünauer, SVN Vishwanathan, Alex J Smola, and Hans-Peter Kriegel. Protein function prediction via graph kernels. Bioinformatics, 21(Suppl 1):i47–i56, 2005.

[8] Esti Yeger-Lotem and Roded Sharan. Human protein interaction networks across tissues and diseases. Frontiers in genetics, 6:257, 2015.

[9] Matthew E Ritchie, Belinda Phipson, D. Wu, Yifang Hu, Charity W Law, Wei Shi, and Gordon K Smyth. limma powers differential expression analyses for rna-sequencing and microarray studies. Nucleic acids research, 43(7):e47–e47, 2015.

[10] Peter Langfelder and Steve Horvath. Wgcna: an r package for weighted correlation network analysis. BMC bioinformatics, 9:1–13, 2008.

[11] Guangyan Zhou, Shuzhao Li, and Jianguo Xia. Network-based approaches for multi-omics integration. Computational methods and data analysis for metabolomics, pages 469–487, 2020.

[12] Angela P Presson, Eric M Sobel, Jeanette C Papp, Charlyn J Suarez, Toni Whistler, Mangalathu S Rajeevan, Suzanne D Vernon, and Steve Horvath. Integrated weighted gene co-expression network analysis with an application to chronic fatigue syndrome. BMC systems biology, 2:1–21, 2008.

[13] Laura L Elo, Henna Jarvenpäa, Matej Orešič, Riitta Lahesmaa, and Tero Aittokallio. Systematic construction of gene coexpression networks with applications to human t helper cell differentiation process. Bioinformatics, 23(16):2096–2103, 2007.

[14] Nir Friedman, Michal Linial, Iftach Nachman, and Dana Pe’er. Using bayesian networks to analyze expression data. In Proceedings of the fourth annual international conference on Computational molecular biology, pages 127–135, 2000.

[15] Anatole Ghazalpour, Sudheer Doss, Bin Zhang, Susanna Wang, Christopher Plaisier, Ruth Castellanos, Alec Brozell, Eric E Schadt, Thomas A Drake, Aldons J Lusis, et al. Integrating genetic and network analysis to characterize genes related to mouse weight. PLoS genetics, 2(8):e130, 2006.

[16] Jesper Tegner, MK Stephen Yeung, Jeff Hasty, and James J Collins. Reverse engineering gene networks: integrating genetic perturbations with dynamical modeling. Proceedings of the National Academy of Sciences, 100(10):5944–5949, 2003.

[17] Albert-Laszlo Barabasi and Zoltan N Oltvai. Network biology: understanding the cell’s functional organization. Nature reviews genetics, 5(2):101–113, 2004.

[18] Dokyoon Kim, Je-Gun Joung, Kyung-Ah Sohn, Hyunjung Shin, Yu Rang Park, Marylyn D Ritchie, and Ju Han Kim. Knowledge boosting: a graph-based integration approach with multi-omics data and genomic knowledge for cancer clinical outcome prediction. Journal of the American Medical Informatics Association, 22(1):109–120, 2015.

[19] Hiromi WL Koh, Damian Fermin, Christine Vogel, Kwok Pui Choi, Rob M Ewing, and Hyungwon Choi. iomicspass: network-based integration of multiomics data for predictive subnetwork discovery. NPJ systems biology and applications, 5(1):22, 2019.

[20] Francis E Agamah, Jumamurat R Bayjanov, Anna Niehues, Kelechi F Njoku, Michelle Skelton, Gaston K Mazandu, Thomas HA Ederveen, Nicola Mulder, Emile R Chimusa, and Peter ACt Hoen. Computational approaches for network-based integrative multi-omics analysis. Frontiers in Molecular Biosciences, 9:1214, 2022.

[21] Martin Oti, Berend Snel, Martijn A Huynen, and Han G Brunner. Predicting disease genes using protein–protein interactions. Journal of medical genetics, 43(8):691–698, 2006.

[22] Lude Franke, Harm Van Bakel, Like Fokkens, Edwin D De Jong, Michael Egmont-Petersen, and Cisca Wijmenga. Reconstruction of a functional human gene network, with an application for prioritizing positional candidate genes. The American Journal of Human Genetics, 78(6):1011–1025, 2006.

[23] Albert-Lászl Barabasi. Scale-free networks: a decade and beyond. science, 325(5939):412–413, 2009.

[24] Charles J Vaske, Stephen C Benz, J Zachary Sanborn, Dent Earl, Christopher Szeto, Jingchun Zhu, David Haussler, and Joshua M Stuart. Inference of patient-specific pathway activities from multidimensional cancer genomics data using paradigm. Bioinformatics, 26(12):i237–i245, 2010.

[25] Bo Wang, Aziz M Mezlini, Feyyaz Demir, Marc Fiume, Zhuowen Tu, Michael Brudno, Benjamin Haibe-Kains, and Anna Goldenberg. Similarity network fusion for aggregating data types on a genomic scale. Nature methods, 11(3):333–337, 2014.

[26] Conghao Wang, Wu Lue, Rama Kaalia, Parvin Kumar, and Jagath C Rajapakse. Network-based integration of multi-omics data for clinical outcome prediction in neuroblastoma. Scientific Reports, 12(1):15425, 2022.

[27] John N Weinstein, Eric A Collisson, Gordon B Mills, Kenna R Shaw, Brad A Ozenberger, Kyle Ellrott, Ilya Shmulevich, Chris Sander, and Joshua M Stuart. The cancer genome atlas pan-cancer analysis project. Nature genetics, 45(10):1113–1120, 2013.

[28] Frank J Massey Jr. The kolmogorov-smirnov test for goodness of fit. Journal of the American statistical Association, 46(253):68–78, 1951.

[29] Khosrow Dehnad. Density estimation for statistics and data analysis, 1987.

[30] F. Pedregosa, G. Varoquaux, A. Gramfort, V. Michel, B. Thirion, O. Grisel, M. Blondel, P. Prettenhofer, R. Weiss, V. Dubourg, J. Vanderplas, A. Passos, D. Cournapeau, M. Brucher, M. Perrot, and E. Duchesnay. Scikit-learn: Machine learning in Python. Journal of Machine Learning Research, 12:2825–2830, 2011.

[31] David W Scott. Multivariate density estimation: theory, practice, and visualization. John Wiley & Sons, 2015.

[32] Michiel JL De Hoon, Ryan J Taft, Takehiro Hashimoto, Mutsumi Kanamori-Katayama, Hideya Kawaji, Mitsuoki Kawano, Mami Kishima, Timo Lassmann, Geoffrey J Faulkner, John S Mattick, et al. Crossmapping and the identification of editing sites in mature micrornas in high-throughput sequencing libraries. Genome research, 20(2):257–264, 2010.

[33] Mark D Robinson, Davis J McCarthy, and Gordon K Smyth. edger: a bioconductor package for differential expression analysis of digital gene expression data. bioinformatics, 26(1):139–140, 2010.

[34] Antonio Colaprico, Tiago C Silva, Catharina Olsen, Luciano Garofano, Claudia Cava, Davide Garolini, Thais S Sabedot, Tathiane M Malta, Stefano M Pagnotta, Isabella Castiglioni, et al. Tcgabiolinks: an r/bioconductor package for integrative analysis of tcga data. Nucleic acids research, 44(8):e71–e71, 2016.

[35] Charles R. Harris, K. Jarrod Millman, Stéfan J. van der Walt, Ralf Gommers, Pauli Virtanen, David Cournapeau, Eric Wieser, Julian Taylor, Sebastian Berg, Nathaniel J. Smith, Robert Kern, Matti Picus, Stephan Hoyer, Marten H. van Kerkwijk, Matthew Brett, Allan Haldane, Jaime Fernández del Río, Mark Wiebe, Pearu Peterson, Pierre Gérard-Marchant, Kevin Sheppard, Tyler Reddy, Warren Weckesser, Hameer Abbasi, Christoph Gohlke, and Travis E. Oliphant. Array programming with NumPy. Nature, 585(7825):357–362, September 2020. doi: 10.1038/s41586-020-2649-2. URL https://doi.org/10.1038/s41586-020-2649-2.

[36] Wes McKinney. Data Structures for Statistical Computing in Python. In Stéfan van der Walt and Jarrod Millman, editors, Proceedings of the 9th Python in Science Conference, pages 56–61, 2010. doi: 10.25080/Majora-92bf1922-00a.

[37] Pauli Virtanen, Ralf Gommers, Travis E. Oliphant, Matt Haberland, Tyler Reddy, David Cournapeau, Evgeni Burovski, Pearu Peterson, Warren Weckesser, Jonathan Bright, Stéfan J. van der Walt, Matthew Brett, Joshua Wilson, K. Jarrod Millman, Nikolay Mayorov, Andrew R. J. Nelson, Eric Jones, Robert Kern, Eric Larson,J J Carey, İlhan Polat, Yu Feng, Eric W. Moore, Jake VanderPlas, Denis Laxalde, Josef Perktold, Robert Cimrman, Ian Henriksen, E. A. Quintero, Charles R. Harris, Anne M. Archibald, Antonio H. Ribeiro, Fabian Pedregosa, Paul van Mulbregt, and SciPy 1.0 Contributors. SciPy 1.0: Fundamental Algorithms for Scientific Computing in Python. Nature Methods, 17:261–272, 2020. doi: 10.1038/s41592-019-0686-2.

